# Heterotypic directional motifs contribute to TAD boundary function in *Drosophila*

**DOI:** 10.1101/2025.09.06.674533

**Authors:** Martina Varisco, Gabriel R. Cavalheiro, Rebecca R. Viales, Christoph Schaub, Charles Girardot, Mattia Forneris, Aaron Comeault, Eileen E.M. Furlong

## Abstract

While convergent CTCF binding sites are essential for topologically associating domain (TAD) formation in mammals, how TADs are formed in other species remains unclear. In *Drosophila*, TAD boundaries lack convergent CTCF sites and are occupied by different insulator protein combinations with no obvious pairing rules. To identify sequence features required for boundary pairing, we inserted different endogenous boundaries into two *Drosophila* TADs in both orientations, creating 53 ectopic insertions and measured their ability to form interactions with the endogenous boundaries to their left and right. 70% of inserted boundaries had context- and/or orientation-dependent effects, indicating specificity and directionality encoded in their sequence. Known insulator motifs or promoters could not explain boundary specificity. Instead, we identified heterotypic pairs of directional motifs that predict compatible boundary pairs. Our results indicate that directional motif pairs are a conserved property of boundaries from flies to mammals, but gained complexity in *Drosophila* through heterotypic combinations.

## INTRODUCTION

Chromatin 3D organization contributes to the regulation of multiple processes, including transcription^1,2^, replication^3^ and DNA repair^2^. During interphase, the chromosomes of many species are dynamically folded into a series of Topologically-Associating Domains (TADs), regions that preferentially self-interact through the formation of chromatin loops^4–6^. A gene’s regulatory elements (enhancers) are often, although not always, contained within the same TAD as the target gene^7^, while TADs typically harbor multiple genes and their enhancers^6–9^. While the presence of chromatin domains and their functions appear largely conserved from flies to humans, how TADs are formed across species remains debated.

Mammalian TADs are formed through cohesin-mediated loop extrusion, which is stabilized through interactions of the insulator protein CTCF^10–14^. As a consequence, mammalian TAD boundaries typically have CTCF bound to convergent binding sites at the loop anchors, leading to focal high frequency interactions at the corners of TADs due to stalling of extruding cohesin loops. In contrast, the mechanism of TAD formation in *Drosophila* remains unclear. The general absence of CTCF::CTCF-bound foci at the loop anchors of *Drosophila* TADs has led to the suggestion that there are no cohesin-dependent structural loops^15^. Different mechanisms for *Drosophila* domain formation have been proposed, including being driven by compartmental interactions^16^, transcription^17^, insulator-pairing^18,19^, or through different insulators (or other compatible proteins) blocking an as yet unknown loop extruder (similar to CTCF in mammals).

Compartments correspond to transcriptionally active and inactive regions, and appear as a checkerboard pattern along chromosomes in Hi-C contact matrices in both *Drosophila*^5^ and mammals^4,20^. Compartments therefore reflect transcription, and as a correlate, transcription and histone marks are among the best predictors of Hi-C contacts in *Drosophila*, *C. elegans* and *A. thaliana*^16,17,21^. Moreover, TAD boundaries are enriched in highly transcribed genes and often contain active promoters in different species, from *Drosophila*^5,17,22,23^, to mouse^4,24^, human^4,11^ and *A. thaliana*^25^. Mammalian boundaries, for example, often occur near highly transcribed housekeeping genes and tRNA^4,11,26^, while 77% of *Drosophila* boundaries overlap at least one promoter^23^. These observations suggest a role for transcription or an active promoter in boundary or insulation function. However, the magnitude varies depending on the locus: inhibition of transcription^16,27,28^ or deletion of a strong promoter at a boundary^29^ leads to a weakening of TAD boundary insulation but not a complete loss, and in many cases has very minor effects^16,27,28,30^.

Insulator-pairing has been proposed based on the genetic dissection of four well-characterised *Drosophila* insulators in transgenic assays (scs–scs, scs’–scs’, 1A2–1A2 and Wari–Wari)^31^. Although it remains unclear if TAD boundaries and these classic insulators represent the same biological entity, they share common properties – insulator elements typically flank a gene and its enhancers, representing small chromatin domains, and are bound by insulator proteins similar to TAD boundaries, but typically do not contain promoters. In transgenic assays, these insulator elements can only interact (or pair) in one orientation, indicating directionality, and mixed pairs could not interact, indicating some specificity^31^. In the bithorax cluster, which contains three genes whose regulatory elements are each bordered by an insulator element, the MCP insulator has orientation dependent effects^32^. Similarly, the function of the Fab-7 insulator element could only be partially replaced by another insulator, indicating specificity in its interactions^33,34^. Directional pairing interactions was also recently shown between the homie insulator elements at the *eve* locus^18,19^. As these classic insulators pairs correspond to borders of chromatin domains, it suggests that orientation dependent boundary function could be much more widespread genome-wide. Such directionality must be encoded in the DNA sequence. However, although individual insulator proteins were identified that are essential for each of the insulators’ function described above (e.g. Zw5, CTCF, Su(Hw))^35,36^, no directional motif arrangements such as convergent CTCF::CTCF sites were detected in any example to date that could explain their directionality or specificity.

In addition to CTCF, *Drosophila* has many other insulator proteins (e.g. Beaf-32, Su(Hw), Pita, M1BP, Zw5, CP190 and others^37^), that are bound in different combinations at endogenous TAD boundaries with no obvious motif arrangement^5,23,29,38–44^. Removal of any one of these factors (such as CTCF) alone reduces insulation at a subset (∼ 10-20%) of boundaries^23,29,42–44^. However, there is currently no single factor or mechanism, which upon depletion leads to the disruption of the vast majority of TAD-level organization in *Drosophila,* in contrast to CTCF depletion in mammals^6^. Redundancy between these insulator proteins has been proposed, although recent studies indicate that it is not simple redundancy where any insulator protein can replace another^29,40^, suggesting some context specificity. Similar to CTCF::CTCF in mammals, interactions of compatible proteins bound to the two genomic regions that will form the loop anchors defining a boundary are presumably required. However, it is currently not clear what rules underly such pairing interactions (and directionality) between boundary elements – for example, what are the motifs required and their orientation?

Here, we set out to determine the sequence rules that shape TAD boundary formation in *Drosophila*, asking if different boundaries are interchangeable, and therefore sufficient to create domains, and – if not – what is the regulatory logic underlying boundary interactions to form a TAD. We used a gain-of-function approach, by inserting endogenous TAD boundaries with different properties into two ectopic locations (the middle of two TADs). Overall, we generated 53 new genetic lines with individual boundary insertions in either orientation into two acceptor TADs, providing interaction data for 106 ectopic-boundary::endogenous-boundary pairs. Assessing their impact using two orthogonal approaches (DNA FISH and Tiled-Capture Hi-C), revealed that the majority of boundaries can function in an ectopic location – indicating pairing with the endogenous boundary to their left or right. While some boundaries could function in all scenarios, the majority displayed some level of specificity, either working only in a specific context (one of the two acceptor TADs), or specific orientation, or a combination of both, indicating specificty in the boundary interactions (i.e. pairing specificity). The occupancy of known insulator proteins or the presence of a promoter was not sufficient to explain the boundaries’ directionality or specificity. Searching for conserved motifs within these compatible boundary::boundary pairs across different *Drosophila* species, identified heterotypic directional motif pairs that explains almost 80% of the tested new boundary::boundary pairs. This includes many motifs for known transcription factors (TFs) that have not been implicated in TAD boundary function to date, in addition to new motifs for unknown factors. We confirm the requirement for some of those TFs by loss-of-function experiments, resulting in chromatin conformation changes at the tested locus. The complexity of motif pairs explains why it was not possible to discern these rules through genome-wide DNA sequence alone. Our extensive functional data uncovered directional motif pairs as the defining property of functional boundary pairs, which appears to be a conserved principle from flies to humans. In *Drosophila,* this consists of directional pairs of heterotypic motifs.

## RESULTS

### An *in vivo* gain-of-function approach for boundary activity

To determine which features contribute to forming a boundary pair in *Drosophila*, we assessed which boundaries can form interactions when placed in an ectopic genomic location (Fig. 1A). TAD boundaries with different properties were inserted into the middle of two TADs (‘acceptor’ TADs) in both orientations (in as far as possible) and their ability to form a new boundary measured by DNA FISH and Tiled-Capture Hi-C (Fig 1A).

**Fig. 1:**
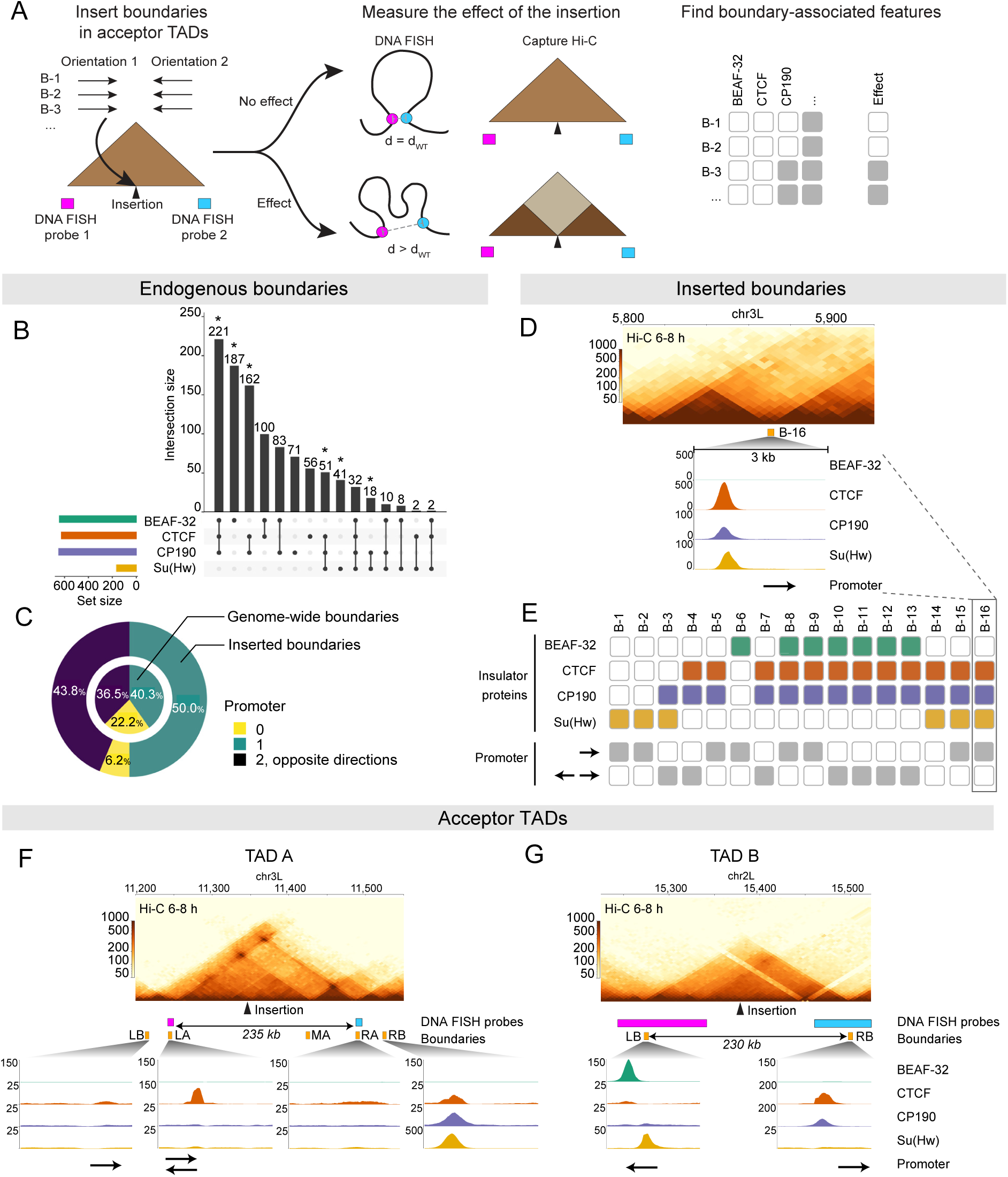
Assessing the rules for TAD boundary formation in *Drosophila* using an endogenous gain-of-function approach. **(A)** Overview of the experimental design. **(B)** UpSet plot showing the genome-wide co-occupancy (ChIP-seq) of 4 insulator proteins, BEAF-32, CTCF, CP190, Su(Hw), at TAD boundaries at 6-8h of embryogenesis. Combinations marked with an asterisk were included in the set of tested inserted boundaries. **(C)** Proportion of genome-wide (inner) and inserted boundaries (outer) that contain a promoter. The number and orientation are indicated. **(D)** Example of an inserted boundary, B-16. Hi-C contact matrix and ChIP-seq of insulator proteins in 6-8 h embryos is shown, with the promoter location and orientation indicated by the arrow. **(E)** Table showing the composition of all inserted boundaries. Filled squares indicate the occupancy of the indicated insulator proteins at the boundary (compare B-16 to panel D). The presence and direction of promoters is indicated in grey. **(F, G)** 3D structure and composition of boundaries of the two acceptor TADs - TAD A (*scyl*-*chrb*) **(F)** and TAD B (*sna-wor-esg*) **(G)**. *Upper*: Hi-C contact matrix in 6-8 h embryos, with the location of the RMCE insertion site (black arrow head), DNA FISH probes (coloured rectangles) indicated underneath. *Lower*: zoom in of the boundary region, showing ChIP-seq insulator binding and promoter presence and orientation (arrows). LB = Left boundary, RB = Right boundary. Additionally, TAD A has three intra-TAD loop anchors (left = LA, middle = MA and right = RA).

Most *Drosophila* TAD boundaries are occupied by different combinations of insulator proteins, as we observed here by ChIP-seq during mid-embryogenesis for four major insulators (Fig. 1B, Table S1), and as previously reported at other embryonic stages^29,45,46^, and in cell lines^23,38,39^. In addition, TAD boundaries often contain active promoters in different species. Of the 1,567 *Drosophila* TAD boundaries at mid-embryogenesis (6-8h, Methods), 77.8% contain a promoter in line with previous reports in a *Drosophila* cell line^23^, 51.8% of which contain a single promoter while 48.2% contain divergent promoters (Fig 1C). To cover this heterogenity, we selected 16 endogenous boundaries that span a range of occupancy patterns for known insulators, including examples of common and more infrequent combinations (Fig. 1B, D, E, Fig. S1), in addition to either a single (50%, n=8), divergent (43.8%, n=7) or no (6.2%, n=1) promoter (Fig. 1C). The latter allowed us to assess if the presence of a promoter is strictly required for boundary function. We included multiple examples with the same insulator binding and promoter properties to examine the sufficiency of each combination (Fig. 1D, E). Each of these 2-3kb endogenous boundary regions were inserted into the same site in the middle (roughly) of the two acceptor TADs by recombination-mediated cassette exchange (RMCE)^47^ (Fig. 1A, Methods).

The two acceptor TADs, *scyl-chrb* locus (TAD A) and *sna-esg-wor* locus (TAD B), were selected as they differ in their internal structure and the composition of their boundaries. TAD A (310 kb) has an internal three-way interaction between a left anchor (LA) upstream of *scylla* (*scyl)*, a middle intergenic anchor (MA) and a right anchor (RA) at *charybde* (*chrb*), all with CTCF binding, leading to two sub-TADs (Fig. 1F). Such a complex 3D arrangement could protect the TAD from disruption by the inserted boundary, or illustrate a gradual or complex pattern of insertion-mediated topological alterations. TAD B (230 kb) is a more archetypical *Drosophila* TAD, with no high-frequency loops and more uniform interaction frequency throughout the TAD (Fig 1G). The boundaries of TAD A and B also have different insulator protein binding and promoter composition, which further increases the diversity of boundary combinations that could be assessed for compatibility (Fig. 1F-G).

The 16 selected boundaries were inserted into TAD A and B using PhiC31 integrase^47^, which can yield insertions in either orientation as desired (Fig. 1F,G). Out of 64 possible insertions (16 boundaries in 2 acceptor TADs in 2 orientations), we obtained 53 lines (∼83%), after several injection rounds: 27 in TAD A (12 boundaries in both orientations, 3 in one) and 26 in TAD B (11 boundaries in both orientations, 4 in one).

### Many inserted boundaries have context- or orientation-specific effects

If proteins bound to an inserted boundary can form interactions with proteins at the endogenous boundary to its left or right, and thus change TAD structure, it should affect the distance between the acceptor TAD’s endogenous boundaries (Fig. 1A). To evaluate this, we used DNA FISH to measure the pair-wise distance between the acceptor TAD boundaries in embryos from each of the 53 insertion lines using probes centered at the loop anchors to the left and right (LA and RA) in TAD A (235 kb distance), and at left and right boundary (LB and RB) in TAD B (∼230 kb distance) (Fig 1F, G). The distances were measured in at least 200 nuclei, in the same region from at least three stage 11 embryos (the stage overlapping the 6-8h Hi-C^48^ and insulator occupancy data) (Fig. 2A, S2).

**Fig. 2:**
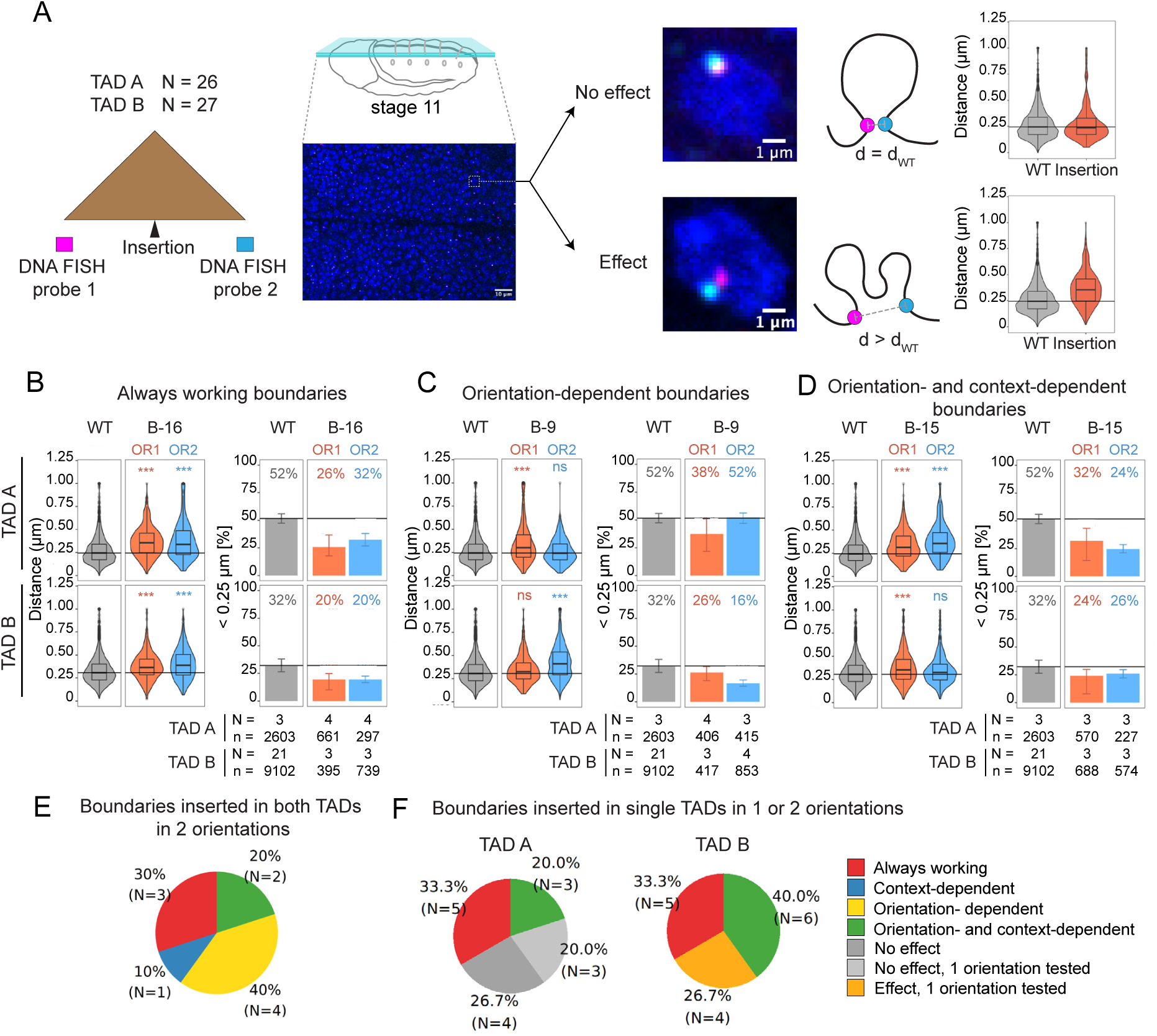
Most boundaries have an effect in the acceptor TAD, often dependent on the orientation and/or context. **(A)** Overview of the DNA FISH measurements. The point-to-point distance between the endogenous DNA boundaries was assessed using FISH probes spanning the boundaries (as indicated) in embryos from each insertion. The schematic embryo (middle) indicates the location (central portion of the dorsal epidermis) where all measurements were taken from, in stage 11 (6-8 h) embryos. Right: Representative nuclei for an insertion that had no effect (upper, overlapping spots) or had an effect (lower, increased distance between the probes). Violin plots show the distribution of distances between the FISH probes (TAD boundaries) in wildtype (grey) or insertion (red) embryos. **(B)** Example of a boundary that works in all 4 conditions (both orientations, both locations) – called always working (B-16). Violin plot (left) showing the distance (uM) and bar plot (right) showing the percentage of nuclei with DNA FISH probes at distances < 250 nm along (median and standard deviation indicated) in TAD A (top) and TAD B (bottom), inserted in orientation (OR) 1 (red) or 2 (blue). Adjusted P value of Kolmogorov-Smirnov test: * p < 0.01, ** p < 0.001, *** p < 0.0001. N = number of embryos, n = number of spots measured, indicated below. **(C)** Same as (B), showing an orientation-dependent boundary (B-9). **(D)** Same as (B), showing an orientation- and context-dependent boundary, B-15, which only works in both orientations in TAD A, but only in OR1 in TAD B. **(E)** Overview of the effects of boundaries inserted in both TADs and both orientations. **(F)** Overview of the effects of all boundaries inserted in at least 1 orientation in TAD A (left) and B (right).

In TAD A, the median distance between the DNA FISH probes is 248 nm in wild type embryos, with 52% of alleles being < 250 nm distance, i.e. below the diffraction limit (Fig 2B, S2). Half of the 27 boundary insertions caused a significant shift in this distance distribution, suggestive of a change in chromatin topology: 5 boundaries had an effect in both orientations (10 insertions), while 3 were orientation-dependent (Fig. 2B-D, 2F, S2). The effect was always an increase in the distance between the endogenous TAD boundaries, indicating a loss or reduction of interactions of the endogenous anchors of TAD A with each other, presumably due to new interactions with the inserted boundary (which we confirm below). Interestingly, the insertion of 7 boundaries had no effect, 4 of which were inserted in both orientations (Fig. 2F), suggesting that they cannot interact with either the left or right endogenous acceptor TAD A boundaries.

In TAD B, the median distance between the DNA FISH probes in wild type embryos is 330 nm, with 32% of alleles being < 250 nm (Fig. 2B, S2). The majority of boundary insertions (80% = 21/26) had a significant effect, a higher proportion than in TAD A. Among the 11 boundaries inserted in both orientations (22 insertions), 6 worked in both orientations (12 insertions), while 5 had orientation specific effects (Fig. 2F). All boundaries inserted in a single orientation (4 insertions) also had an effect. Similar to TAD A, the majority of insertions led to an increase in the distance between the endogenous boundaries, suggesting that the inserted element can interact with one or both of the endogenous TAD’s boundaries (Fig. S2). However, there are three interesting examples in TAD B where the insertion led to a decrease in the distance between probes (Fig. S2, black arrow heads), hinting at TAD compaction, potentially through the formation of a three-way interaction between the endogenous boundaries and the inserted boundary, or by the stabilization of a smaller DNA loop.

We obtained a complete set of insertions for ten boundaries, leading to 40 boundary insertion *Drosophila* lines:10 boundaries x 2 orientations x 2 acceptor TADs. This enabled a more direct comparison, and revealed interesting cases of context and orientation dependent specificity. Three boundaries function regardless of the context (acceptor TAD) or orientation, disrupting TAD A and B in both orientations (12 insertions). In contrast, one boundary was context-dependent (disrupting TAD B in both orientations, but had no impact on TAD A), two were orientation-dependent (disrupting both TADs in a single orientation), while four were orientation- and context-dependent (disrupting only one TAD, in a single orientation) (Fig. 2E). These results indicate that many *Drosophila* TAD boundaries have context-specificity, suggesting specific pairing rules encoded in their sequence. Moreover, the orientation dependency of the inserted genomic regions suggests directionality in the motifs that they contain. Together, this indicates that the directionality and context specificity observed in transgenic assays with the five well-studied ‘classic’ *Drosophila* insulators also holds for TAD boundaries, and more globally in an endogenous context.

### Inserted boundaries can lead to diverse changes to chromatin topology

To determine the topological changes induced by the boundary insertions, and to have an orthogonal measurement to DNA FISH, we developed a multiplexed Tiled-Capture Hi-C protocol detailed in the Methods. We performed Tiled-Capture Hi-C on 4-20 h embryos (after egg laying) from 38 lines – which includes a wild-type reference and 37 insertions composed of 24 insertions in TAD A and 13 insertions in TAD B. As TAD A contains multiple potential regions that could interact with the inserted boundary (3 loop anchors and 2 TAD boundaries), the Tiled-Capture Hi-C was particularly important to determine which elements engage in interactions with the inserted elements. We therefore examined all insertions with an effect by DNA FISH in TAD A (13 working insertions, including the 3 orientation-dependent boundaries), and 11 lines with no apparent effect. TAD B has a more simple structure, with a single boundary at either side and no internal loops. Given this, we selected 11 working insertions and 2 with no effect by FISH, prioritizing boundaries that were inserted in both TAD A and B, and have different insulator protein occupancy. From the selected 11 working boundaries, eight had an effect in both orientations (3 of which led to compaction), and 3 were orientation-dependent (Fig. S2).

Insertions that can act as a boundary in an ectopic location were expected to split the TAD into one or two sub-domains, by forming interactions with the boundary to their left or right. While we observed this, the insertions led to a much broader range of topological effects, especially in TAD A (Fig. 3 and Fig. S3-6). This includes a local loss of contacts (e.g. B-3 OR1, Fig. 3A), the creation of sub-TADs while still partly preserving the integrity of the original acceptor TAD (B-3 OR2, Fig. 3A), combined with either a strengthening (e.g. B-3 OR2, Fig. 3A), weakening (e.g. B-14 OR1, Fig. 3A) or having no effect on chromatin loops (e.g. B-13, OR1/2, Fig. S3), and in the most extreme cases a complete disruption of both the TAD and loops (e.g. B-14 OR2, Fig. 3A) (Table S2). Notably, boundary pairs that always worked by DNA FISH (defined in Fig. 2B and S2) regardless of their context or orientation were typically associated with larger effects on intra-TAD interactions (sub-TADs or TAD disruption) and at loops (e.g. B-14, Fig. 3B, S3), compared to insertions with orientation-specific effects, which tended to have more subtle changes (B-9 OR1 and B-4 OR2, Fig. 3B).

**Fig. 3:**
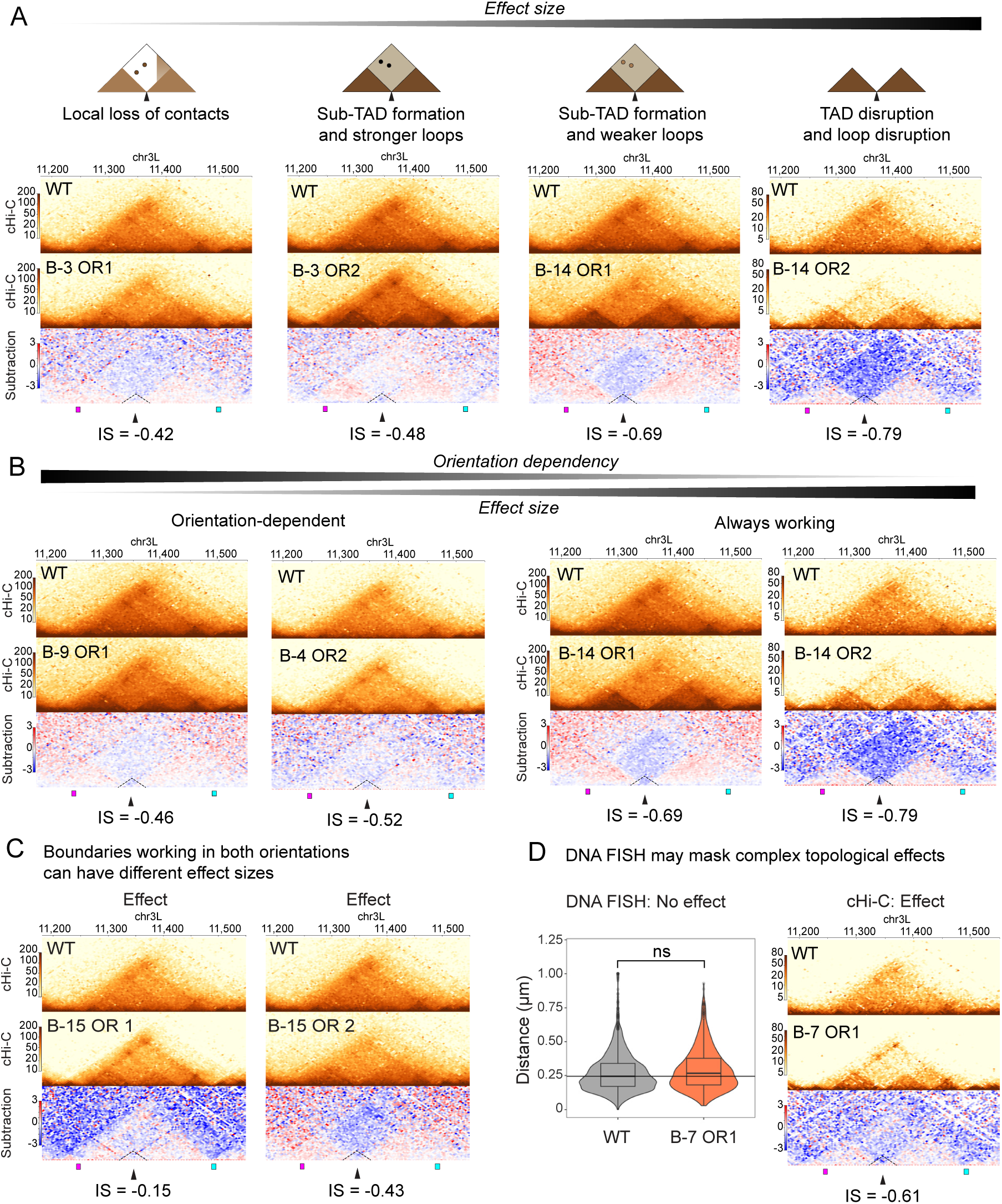
Tiled Capture Hi-C uncovers the changes in TAD structure and interacting regions. **(A)** Boundaries with an effect in TAD A can disrupt the TAD to different extents, from a reduction of local interactions (left), to sub-TAD formation and changes in loop strength (middle), to complete spitting of the TAD to two sub-TADs (right). Contact matrices of Tiled-Capture Hi-C (cHi-C) in wild-type (WT) embryos (top), in the boundary insertion line (middle) and the subtraction map (bottom), and relative insulation score (IS) below. For each WT-insertion pair, cHi-C maps were down-sampled to the same number of reads for a more quantitative comparison. The boundary number and orientation are indicated. The area close to the insertion locus is marked with a dotted triangle. **(B)** Always working boundaries tend to have a larger effect than orientation-dependent boundaries. cHi-C and subtraction maps as above in orientation-dependent boundaries (B-9 and B-4) (left) and in an always working boundary insertion (B-14) (right) in TAD A. **(C)** Example of an always working boundary that has different effect sizes depending on its orientation (B-15). cHi-C and subtraction map as in (B). **(D)** Example of a boundary (B-7) with no detectable effect by DNA FISH, but a significant impact by cHi-C. Violin plot (left) of DNA FISH probe distances in WT (grey) and boundary insertion (red) embryos and cHi-C matric (right), showing sub-TAD formation and loop strengthening, which is likely masking the detection of changes by DNA FISH.

Interestingly, boundaries with an effect in both orientations in TAD A by DNA FISH appear to have different mechanisms. To obtain a higher resolution, we performed virtual 4C on the normalized Tiled-Capture Hi-C of the matched wildtype (WT) and insertion lines, using the loop anchors and boundaries as viewpoints. By comparing the interaction frequencies across the TADs in the two conditions (WT and insertion), we observed interesting differences (Fig. S3-6): for instance, boundary B-15 created sub-TADs and fuzzier stronger loops in OR1, but led to a general decrease in intra-TAD interactions and loop strength in OR2 (Fig. 3C, S6). From the 11 selected boundaries with no apparent effect in TAD A by DNA FISH, the virtual 4C revealed interesting effects for two. B-7 in OR1 resulted in both a loss of intra-TAD contacts and a strengthening of the loops (Fig. 3D). This combination led to no significant difference in the distance between the two endogenous TAD boundaries by DNA FISH. The second insertion (B-3 OR1) had a small change in the frequency of interactions around the point of insertion, but it was very minor. In all other cases, the results from DNA FISH were confirmed by capture Hi-C (Table S2).

In TAD B, the majority of working boundaries by DNA FISH led to an increase in the distance between the endogenous boundaries (18/21 insertions with an effect). Tiled-Capture Hi-C revealed that these insertions had a more uniform effect on topology (compared to TAD A), with lower interaction frequencies at both the borders and within the TADs, resulting in less defined boundaries, lower intra-TAD interactions, with little obvious TAD splitting (e.g. B-1 OR1, Fig. S7). Only few insertions created fuzzy sub-TADs (B-11 OR2, B-16 OR2, Fig. S7). In contrast, the three boundary insertions that reduced the distance between the endogenous boundaries by FISH had minor changes (by Capture Hi-C) within the TAD and between its boundaries, indicating that such changes may be more easily detectable by imaging methods. Interestingly, boundary B-2 led to compaction when inserted in OR1, but decompaction (or a greater distance between the endogenous boundaries) in OR2 (Fig. S6, S7).

Taken together, the Tiled-Capture Hi-C largely aligns with the DNA FISH results (92% (34/37) of cases), while the measurements of interactions across the entire TAD uncovered the underlying changes in topological structure. This revealed that boundary insertions can change chromatin topology in different ways: altering TAD compaction as a whole, splitting a TAD into sub-TADs while also maintaining the TAD structure, generating two separated TADs, or even affecting its intra-TAD loops. Moreover, it demonstrates that not only TAD decompaction, but also compaction is possible, as shown for three insertions in TAD B. Although some boundaries always worked regardless of their context or orientation (Fig. S3), the majority had context (Fig S4), orientation (Fig. S5), or both context & orientation (Fig. S6) effects, indicating extensive specificity in their ability to form a functional interacting pairs.

### Known insulator protein occupancy and promoter presence are not sufficient to explain boundary directionality and context specificity

We next asked can insulator protein binding for the known factors or the presence of a promoter explain which boundaries can function (establish contacts) in the ectopic locations? Boundary strength was previously shown to correlate with the number and identity of co-bound insulator proteins^29,39^. To assess this within an ectopic context, we grouped the inserted boundaries based on their occupancy (the combination of insulator proteins bound to them), and for each combination counted the number of inserted boundaries that worked or did not work in the ectopic location. The combination of the 3 insulator proteins CTCF + CP190 + Su(Hw) was most strongly linked to working boundaries by DNA FISH, with 6/6 insertions having an effect in TAD A and 5/6 in TAD B (Fig. 4A, B), however they could not explain the directionality or specificity observed. For example, one CTCF + CP190 + Su(Hw) insertion (B-15 in TAD B) was orientation dependent, but the others (B-14, B-16, and B-15 in TAD A) were not (Fig. 4A). Another 3-factor combination (CTCF + CP190 + BEAF-32), despite being the most represented genome-wide at endogenous boundaries (221 boundaries, Fig 1B), only worked in a subset of cases in TAD A (3/8 tested insertions), indicating that the number of insulator proteins does not define functional boundaries. This is also evident from boundaries occupied by two insulator proteins, e.g. CTCF + CP190 had an effect in 4/5 cases in TAD B but only in 3/6 in TAD A, which is a similar efficiency to CTCF + CP190 + BEAF-32 (Fig 4A, B). This indicates that the likelihood of a boundary retaining its function in an ectopic context is influenced by which insulator proteins are co-bound. However, our data does not support a simple additive model where boundaries co-bound by more insulator proteins have progressively stronger activity, and rather highlights context-dependent specificity (Fig. 4B).

**Fig. 4:**
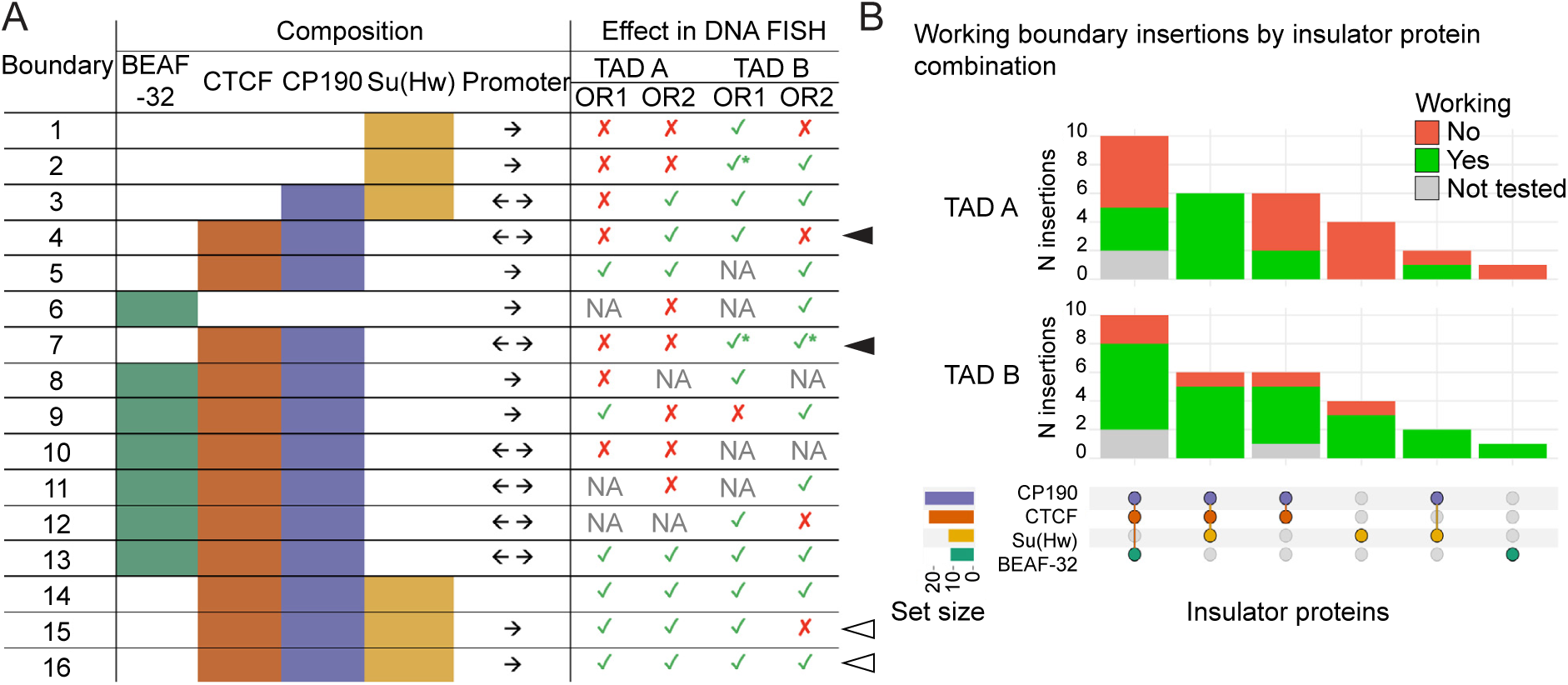
The binding of the most prominent insulator proteins or promoter presence do not explain working versus non-working boundary pairs. **(A)** Overview indicating which boundaries working in which condition (compatible boundary pairs) together with their insulator binding and promoter presence (arrows). Black arrow heads (right) indicate two boundaries with divergent promoters and context-/orientation-specific effect, white arrows indicate two boundaries with a single promoter and orientation-independent effects in at least one TAD. **(B)** Frequency of working (green) and not working (red) insertions in TAD A (top) and B (bottom), grouped by boundary composition. Boundaries not tested in the TAD are indicated in grey.

Promoters and promoter-associated features such as transcription have also been associated with stronger boundary insulation^23,29^. However, our results indicate that the presence of a promoter is not strictly essential for boundary activity, at least in the context of the boundaries tested here. In fact, a boundary without a promoter (B-14) works in both orientations in both TADs (Fig. S2), and divergent promoters are less represented in working boundaries than single promoters: boundaries with divergent promoters worked in 4/13 cases in TAD A and 9/13 cases in TAD B while boundaries with single promoters worked in 7/14 cases in TAD A and 10/13 in TAD B (Fig. 4A, S2). Moreover, the activity of working boundaries bound by the same combination of insulator proteins is not increased by the presence of a divergent promoter. For example, a boundary with CTCF + CP190 and a single promoter appears to work better than a boundary with CTCF + CP190 and a divergent promoter: 3/3 insertions disrupt topology in the former, versus 2/4 in the latter (Fig 4A, S2). Also, the direction of the promoter is not obviously correlated with the direction (orientation dependency) of the boundary insertion: 3/5 orientation-dependent boundaries in TAD A or B have divergent promoters, and 4/10 boundaries working in both orientations have a single promoter (Fig 4A). Therefore, our data does not support a role for promoters in fully determining boundary function or orientation dependency, even though in specific cases promoters can contribute to boundary strength^29^.

Notably, different boundaries with the same known insulator and promoter composition typically do not have the same effect: the CTCF + CP190 + divergent promoter boundaries B-4 and B-7 are orientation-dependent and context-dependent, respectively (Fig. 4A, black arrows). Similarly, the CTCF + CP190 + Su(Hw) single promoter boundaries B-15 and B-16 are orientation- and context-dependent (B-15) or always working (B-16) (Fig. 4A, white arrows). There is therefore no single rule based on insulator protein occupancy or promoter properties that explains the compatibility of working boundary pairs, instead the compatibility appears more context specific, suggesting that additional factors are involved.

### A set of conserved motifs is associated to boundary function

Given the inherent specificity we uncovered for the compatibility of different boundary combinations, we used our functional data to search for DNA motifs and syntax enriched in working boundary pairs compared to non-working pairs. In TAD A, which has two boundaries and loop anchors on the left and right of the insertion, the inserted boundary can have preferential partner(s). The higher resolution of the virtual 4C (Fig. S3-8) could better pinpoint the interacting loop anchors, and identified 79 interacting boundary::boundary pairs or boundary::anchor pairs (an ectopic boundary that can interact with an endogenous TAD boundary or loop anchor), out of 187 potential interactions of the inserted boundaries with the acceptor TAD boundaries (Table S3, examples shown in Fig. S8).

To identify sequence motifs associated with our experimentally tested boundaries (whether they were compatible with making interactions in an ectopic location or not), we used sequence conservation to identify small DNA stretches (potential motifs) of high conservation. TAD boundaries are generally conserved in their function, especially in TADs containing developmental genes, as shown in *Drosophila*^45^ and mice^49^. Indeed, 12 out of 16 of our inserted boundaries in *Drosophila melanogaster* also function as endogenous TAD boundaries in 7 other *Drosophila* species, spanning ∼25 million years of evolution, while an additional two boundaries were not assessed^50^. To identify conserved motifs within these largely functionally conserved boundaries, we performed sequence alignment of the inserted and acceptor TAD boundaries (2-3 kb regions) throughout the *D. melanogaster* clade (containing 19 species, spanning over 20 million years of evolution)^51^, removing coding regions and promoters (Fig. 5A). We focused on sequence stretches with the highest conservation (top 25% conservation score) in DNase Hypersensitive Sites (DHSs) within the boundaries (using *D. melanogaster* DHS data at 6-8h from^52^). Each group of more than 6 consecutive conserved bases within the boundary DHS was considered conserved that could contain a potential motif (Fig. 5A, B). We allowed up to three consecutive bases with lower conservation score within a conserved sequence, in line with the typical flexibility of known motifs scores in Position-Weight Matrices (PWMs). Based on these criteria, we identified 279 conserved sequences within these boundaries, mostly between 6 and 30 nucleotides in length (Fig. 5A, B). Some of these are alternative motif variants that are likely bound by the same factor. These 279 conserved sequences are highly enriched in working boundary pairs compared to non-working boundary pairs (86% (239/279), P< 0.0001).

**Fig. 5:**
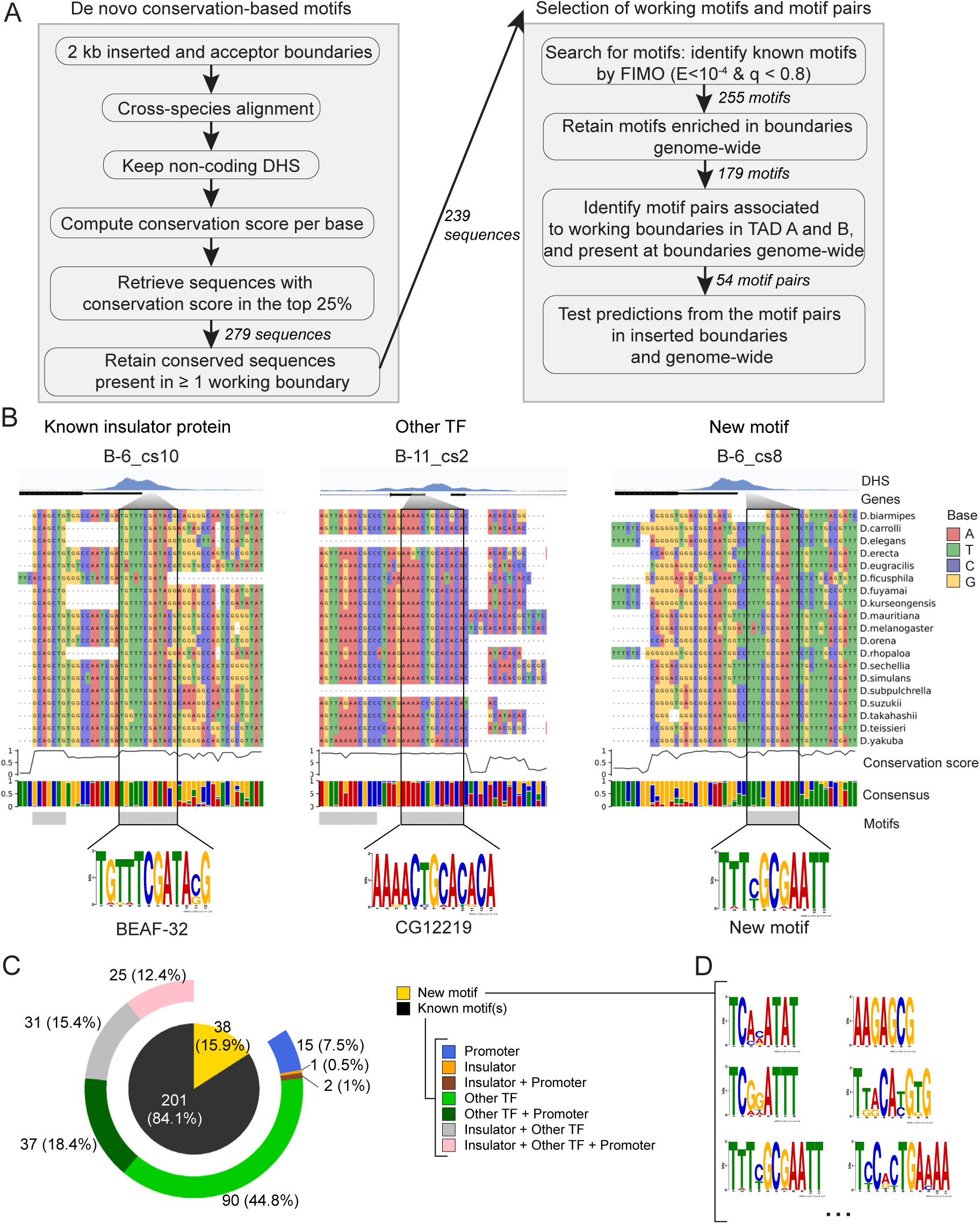
*De novo* identification of conserved motifs enriched at TAD boundaries. **(A)** Computational workflow to identify conserved stretches of sequences (length ≥ 6 bp) in DHS (excluding genes) present in each inserted and acceptor TAD boundary, the majority of which are functionally conserved. Of the 255 identified conserved motifs, 179 are enriched in TAD boundaries genome-wide and were retained. Examining which of those are present at the anchors of working (compatible boundary pairs) versus non-working boundary pairs, identified 54 pairs of 30 motifs in working boundary pairs, one at each boundary. **(B)** Examples of conservation-based motif identification of a known insulator protein (BEAF-32, left), a transcription factor (CG12219, middle) and a new motif with no known corresponding transcription factor (right). From the top, DHS track, genes (horizontal black line), a zoom in of DNA sequence aligned in 19 *Drosophila* species (indicated on the right) in the *D. melanogaster* clade. Bottom: DNA conservation score, the consensus sequence and derived PWM of the conservation-based motif. **(C)** Overview of the composition of the identified conserved sequences: inner circle shows the proportion of conserved sequences with new and known motifs, and the outer ring indicates the type of known motif, promoter (blue), other TFs (green) as indicated. **(D)** Examples of 6 new conserved motifs enriched at TAD boundaries genome-wide.

To identify potential factors that could be binding to theses conserved sequences, we used FIMO^53^ to search for motifs (Methods). Among the 239 conserved sequences present in at least one working boundary pair, 201 (72.1%) contain known motifs (in many cases multiple motifs leading to a total of 255 identified motifs, Methods) attesting to the specificity of the approach (Fig. 5C). The 239 sequences have highly diverse combination of motifs: (i) known insulators alone or combined with promoter motifs or other TF motifs (which together account for ∼30% of the known motifs), (ii) motifs for other TFs alone or in combination with promoter motifs (together account for ∼62%); (iii) promoter motifs alone (7%) (Fig. 5C). 38 conserved sequences do not match any known motif despite the large-collection of curated motifs used, many of which have high sequence information content and are potentially motifs for new factors (Fig. 5B, D).

To assess if these motifs are functionally important for the ability of boundaries to interact, we first determined if the conserved motifs (identified in the 16 tested boundaries) are also enriched at endogenous boundaries, genome-wide. Of the 255 motifs present in the conserved sequences in our test boundaries, 70% (179/255) are also enriched at endogenous TAD boundaries (p < 0.05) (Table S4), including the motifs for all known insulator proteins (with the exception of Trl), promoter motifs (7), motifs for known TFs in the cisBP2.00 database (148), and new motifs (15). We next assessed which of these 179 motifs are associated with working versus non-working boundary pairs. This identified 165 motifs that are exclusive to working (compatible) boundary pairs that function in at least one condition (TAD/orientation), among which 151 are known and 14 are new motifs (Table S5). Again, this includes the motifs for the majority of known insulator proteins, including BEAF-32, CTCF, Trl, M1BP, which are present in boundaries that work in TAD B more often than in TAD A. Pita and some new motifs are present in boundaries that always work in both TADs, and in orientation-dependent boundaries (Table S6). Interestingly, single motifs in inserted boundaries are not sufficient to explain if the boundary can function or not, indicating that additional motifs, most likely in the paired acceptor TAD boundary are likely required to form a working boundary pair.

### Directional heterotypic motif pairs explain boundary function

Given that single motifs cannot explain the pairing compatibility of working boundaries, we searched for pairs of two different motifs within the two compatible interacting boundaries. Remarkably, all 165 conserved motifs associated with working boundaries are always present in pairs, with one member of each pair being present at one end of the 79 working boundary::boundary pairs or boundary::anchor pairs. Given the orientation-dependent effects observed by the majority of inserted boundaries, we reasoned that directional interactions must be driven by directional non-palindromic motifs in the inserted boundary and in one of the endogenous acceptor TAD boundaries or loop anchors. Such a motif architecture would resemble convergent CTCF motifs found at vertebrate boundaries.

To explore the directionality of the conserved motifs, we examined the frequency of each motif pair to be in a convergent, divergent or tandem orientation in working (79) vs non-working (108) boundary pairs (Methods). As more insertions were compatible with the acceptor TAD boundaries in TAD B (80%) compared to TAD A (32%), the frequency of motif pairs in working boundaries will appear higher for TAD B related motifs. To account for this, we required motif pairs to be associated with working boundary pairs in 100% of cases in TAD B, and in > 80% of cases in TAD A. This identified 54 motif pairs associated with working boundaries in a specific directional orientation, of which 15 pairs are convergent, 14 divergent and 25 tandem (Table S7, Fig. 6A).

**Fig. 6:**
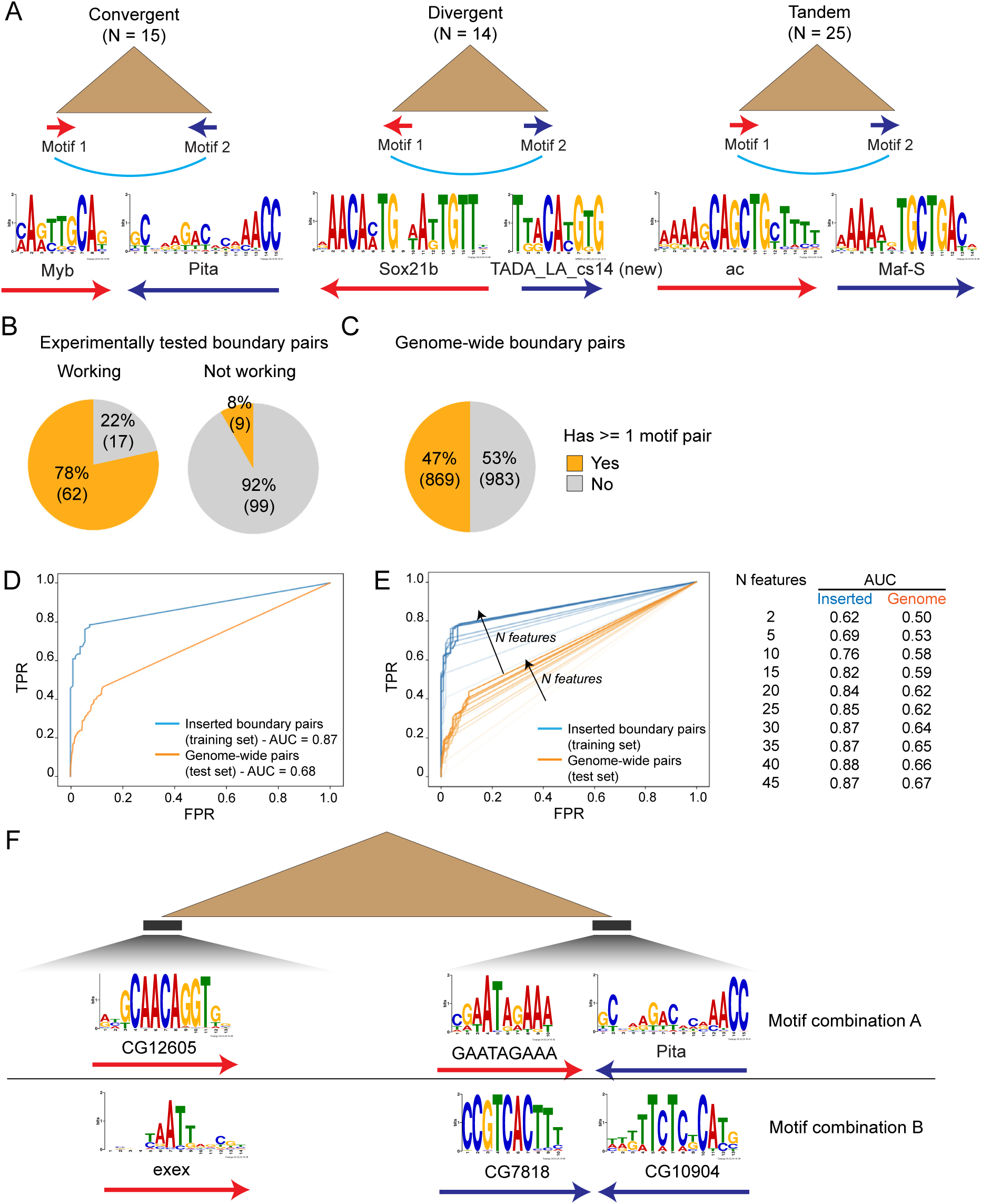
Directional motif pairs are associated with compatible boundary::boundary interactions. **(A)** Examples of convergent, divergent and tandem motif pairs associated with working boundary pairs. **(B)** Percentage of compatible (working) (left) and non compatible (right) tested boundary pairs with at least 1 directional pair of conserved motifs (orange). **(C)** Percentage of genome-wide TAD boundary pairs with at least 1 directional motif pairs (orange). **(D)** ROC curve showing the ability of the identified motif pairs to predict compatible versus non-compatible boundary pairs. For inserted boundaries, working and non-working boundary pairs were used for the positive and negative set, respectively; for the genome, TAD boundaries and pairs of insulator-bound DHSs were used for the positive and negative set, respectively. A logistic model was trained on working and non-working experimental boundary pairs with 5-fold cross-validation, and tested on genome-wide pairs. ROC on the cross-validation test set of inserted boundary pairs (blue) and on genome-wide boundaries (orange). **(E)** ROC curve showing the ability of the identified motifs to distinguish between working boundary pairs in the test set (blue lines) and genome-wide (orange lines), with N motifs from recursive motif elimination. AUC for inserted and genome-wide pairs for different number of features. **(F)** PWMs of motif combinations that involve more than one directional motif pair on each side of a working boundary pair. Motif combination A occurs in 6 instances in the test set and 15 genome-wide, while combination B occurs in 5 instances in the test set and 15 genome-wide.

This set of directional motif pairs explains almost 80% (62/79) of working boundary pairs tested, and are present in only 8% (9/108) of non-working pairs (Fig 6B). 12 of these motif pairs are exclusively associated to boundaries that always worked in both TAD A and B, and may therefore represent a more universal set of motifs (Table S8). 19 motif pairs are associated with orientation-dependent boundaries in at least one TAD. Although our experimental boundary test set used for the motif discovery was relatively small, 47% (869/1852) of all endogenous TAD boundary pairs have at least one of the directional motif pairs (Fig. 6C). We also tested to what extent the discovered motif pairs are predictive of boundary pairs, genome-wide. A logistic regression model was trained on our set of experimentally tested working boundary pairs, using non-working boundaries as background. The trained model could discriminate between working and non-working boundary pairs very well, with a ROC-AUC of 0.87 (Fig. 6D). The trained model was then applied to all boundary-pairs genome-wide, using insulator-bound DHS sequences as a background. This could predict genome-wide boundary pairs reasonably well, with a ROC-AUC score of 0.68 (Fig. 6D), which confirms that this set of motif pairs can discriminate to some extent between true compatible boundary pairs, but also indicates that there are more directional motifs still to be discovered. By feature selection, we could reduce the motif set to 25 motif pairs with negligible loss in prediction accuracy (Fig. 6E).

Among the motif pairs, heterotypic pairs of motifs are the main contributors, in contrast to mammalian boundaries which are dominated by homotypic pairs of CTCF::CTCF motifs. 15 motif pairs involve one known insulator at one boundary with a non-insulator TF at the other boundary. Among the well characterised insulators, the motif for Pita is the most prominent paired with different motifs, followed by Su(Hw) (both C2H2 containing insulator proteins), while BEAF-32 and CTCF were not present in sufficient numbers of working boundaries to be recovered among the top motifs (Table S9). For example, we observed a convergent arrangement of the motif for CG12605, a C2H2 zinc-finger protein (in the forward orientation, left boundary) with Pita (reverse orientation, right boundary), with or without the promoter motif GAATAGAAA – the CG12605 motif has a 62.5% co-occurrence with the Pita motif in this arrangement (Fig. 6F, Table S9). Among the TFs not known to be insulators, the pairs include motifs for ubiquitous (Myb, CG4404, Maf-S) and tissue-specific (Dbx, Vsx, Sox21b, Exex, Bin, Rn, Vnd, Wor, Kr) factors (Table S7). One example of a convergent motif pair involves Exex (a homeodomain TF) and CG10904 (predicted MADF-BESS domain TF) motifs, which co-occurs with CG7818 in 71.4% of cases (Fig. 6F). The motif for Myb is among the most prominent, occurring in 5 motif pairs in working boundary pairs, along with its cofactor E2f2^54^ (Table S7). The Myb motif is more predictive than the motifs of co-localizing insulator proteins, and has a clear directionality preference in both the tested functional boundary pairs and genome-wide, which is not observed for the CTCF motif. Although our identification of Myb was based solely on its presence in directional motif pairs in working boundary pairs, its potential role in TAD formation is supported by its requirement for BEAF-32 binding at a number of sites in wing imaginal discs^55^, and by its physical interaction with CTCF and CP190^56^. Lastly, 4 different directional motif pairs include two new conserved motifs with unknown binding partners (TADA_LA_cs13, TADA_LA_cs14) (Table S7).

These findings suggest that a directional motif-code underlies TAD formation in *Drosophila*, similar to vertebrate boundaries with CTCF::CTCF convergent motifs. In *Drosophila*, the situation is much more complex, involving heterotypic motif pairs in either a convergent or divergent orientation. This opposes previous hypotheses that *Drosophila* TADs are primarily formed by compartmentalization-based mechanisms, and/or as a direct consequence of transcriptional status^16,17^. The heterotypic motif pairs found here provide a foundation for which factors could underlie boundary formation and specificity in *Drosophila*.

### New factors involved in orientation dependent boundary formation

Our functional dissection of what makes a working boundary pair identified motifs for many new potential insulator proteins. To confirm their role in boundary formation, we selected 4 factors whose motifs are present in motif pairs at compatible boundary pairs in TAD A, including Pita as a positive control, and two factors (Bbx, Sox21b) whose motif was not present in TAD A boundaries (but was in TAD B) as negative controls (Fig. 7A). RNAi was used to deplete each factor (both the maternal and zygotic supply) and we assessed the impact on TAD boundary distance by DNA FISH in late stage 5 embryos. If the factor is essential for the boundary to form, the distance between the two FISH probes is expected to increase (similar to the insertion of boundary sequence).

**Fig. 7:**
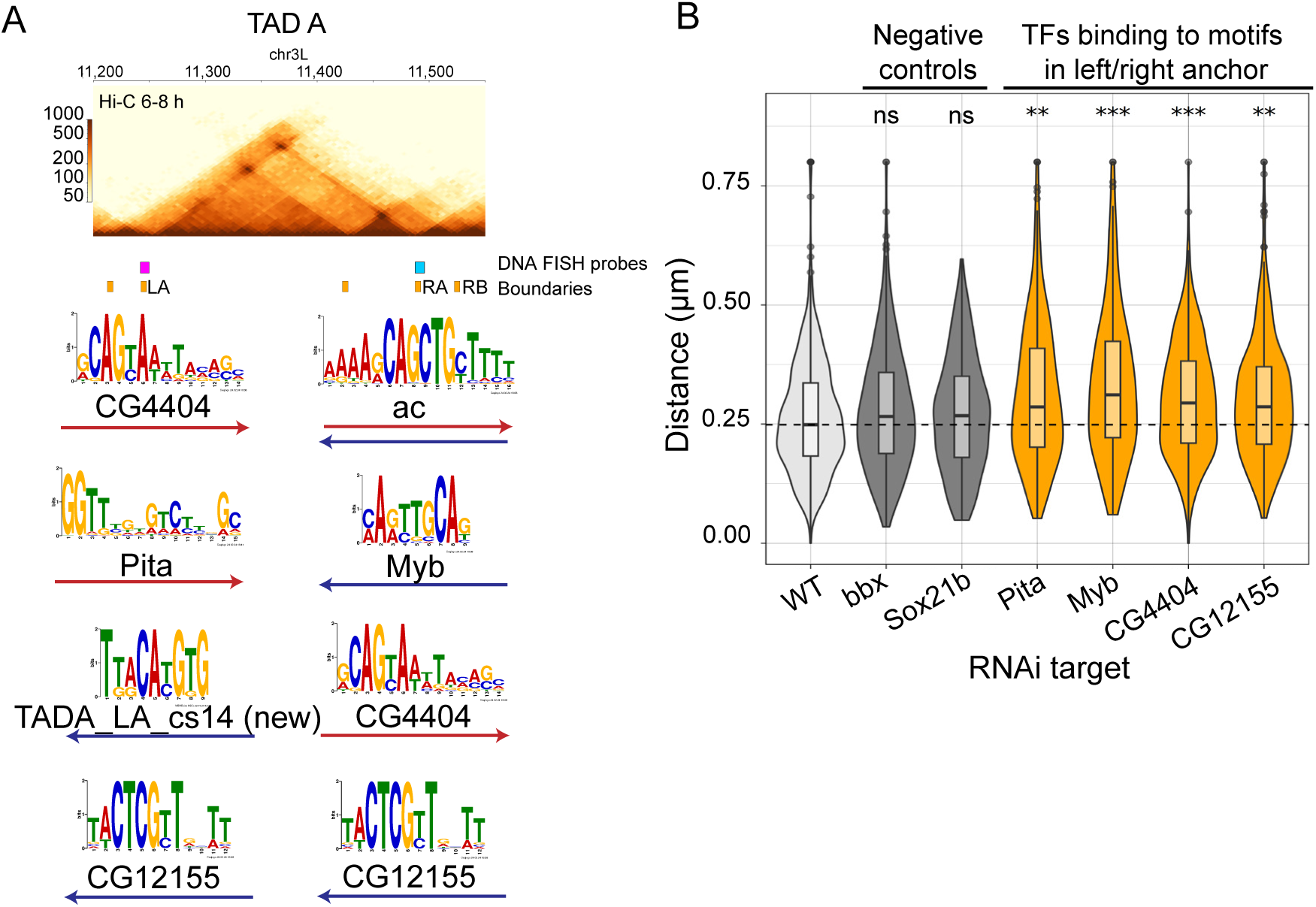
Depletion of TFs bound to conservation-based motif pairs causes topology changes. **(A)** Hi-C map of the TAD used for validation, indicating the discovered motifs pairs, with their directionality indicated. **(B)** Violin plot of the DNA FISH distances upon RNAi depletion of each factor. Negative controls (factors depleted by RNAi whose motif is not present at the TAD’s boundaries) coloured in grey, factors with directional motifs at the boundaries are colored in orange. The point-to-point distance between the endogenous DNA boundaries was assessed using FISH probes spanning the boundaries (as indicated in Fig. 1F) in embryos from each genotype. DNA FISH distances were measured in a similar dorsal region of late 5 embryos, 100 nuclei each, for each genotype. Adjusted P value of Kolmogorov-Smirnov test: * p < 0.01, ** p < 0.001, *** p < 0.0001.

Among the 6 RNAi lines, depletion of Pita and Myb led to over 75% embryonic lethality, recapitulating other studies^57,58^ and confirming the RNAi efficiency. Depletion of the other 4 TFs (Bbx, CG4404, CG12155, Sox21b) led to between 50-70% embryonic lethality. As we have no antibodies for these factors, we cannot assess if the remaining viability is due to the depletion efficiency, or to the biological role of these factors (the function of Bbx, CG4404, CG12155, and Sox21b has never been assessed during embryogenesis to date). However, we assume that the results obtained likely represent partial depletions, with some level of each factor remaining.

Depletion of each TF present at TAD A boundaries led to a significant increase in the distance of the TAD boundaries by DNA FISH (4/4 TFs: Pita, Myb, CG4404 and CG12155), while Bbx and Sox21b depletion, whose motifs are not present, had no effect (Fig. 7B). Our validation shows that Myb, CG4404 and CG12155 – which we found through motif conservation in functional boundary pairs – have a role in boundary formation, alongside the well-known insulator protein Pita. It is interesting to note that, as all 3 pairs of directional motifs are present at the boundaries of this TAD, there appears to be multiple contributors to boundary formation; in fact, 16.8% of the boundary pairs genome-wide have more than 1 motif pair (Table S10). Yet, depletion of only one factor in a pair still has an impact on the TADs formation.

## DISCUSSION

Many studies have identified a number of insulator proteins enriched at *Drosophila* TAD boundaries. However, depletion of any one insulator proteins only leads to changes (reduced insulation) at a subset of boundaries (∼10-20%)^23,29,40,43,44^, suggesting that additional factors or combinations of proteins are involved. By selecting endogenous boundaries and testing their compatibility to form interactions at ectopic sites, we sought to identify the main determinants of compatible boundary pairs. We used endogenous boundaries as they contain the sequence features (motifs beyond known insulator proteins) to form a working boundary pair at their endogenous site and, for the majority of cases, their boundary function is conserved across diverse *Drosophila* species. Our results revealed that many boundaries can retain their function in an ectopic site, by establishing selective interactions with the endogenous boundary to their left or right. Many boundary insertions can split the acceptor TAD into sub-TADs, demonstrating that boundaries have intrinsic activity and TADs are a not a by-product of transcription compartments or chromatin state transitions^5,17^. In contrast, compatible interactions are associated with paired motifs (one at each side) in a preferential orientation, leading to orientation-specific and context-specific boundary function. Known insulator proteins alone are not sufficient to explain the observed directionality and specificity of boundary pairing. We therefore uncovered directional motif pairs, many involving new factors, that contribute to boundary function in *Drosophila*, shifting the paradigm of co-bound insulator proteins to directional sequence determinants analogous to CTCF motifs in mammals. Notably, in *Drosophila*, this predominantly represents heterotypic motif pairs, a fraction of which (17/54 pairs) involve a known insulator protein on one side with a new motif on the other.

### Motif pairing and directionality contribute to boundary function

Specific combinations of insulator proteins are preferentially found at working boundary pairs, being compatible with the acceptor TAD boundaries (e.g CTCF + CP190 + Su(Hw)). However, they are not predictive of context and orientation specificity, indicating that other proteins are involved. Both context- and orientation-dependent boundaries indicate specificity and directionality, which must be imparted by other factors binding to additional motifs within the inserted boundaries. This indicates that it is the sequence properties of the two interacting boundaries that makes them compatible, and not general features of all boundaries, that drive TAD formation. The always working boundaries, independent of orientation and context, may contain multiple motif pairs, thereby maximizing the chance of interactions, or involve TFs that bind to palindromic motifs. However, these were a minority, the majority of working boundaries function in a context- and/or orientation-specific manner.

Orientation-dependent boundaries in particular underscore the role of directional sequence elements, i.e. motifs or promoters in the sequence, that facilitate specific boundary-boundary interactions. Directionality has been observed for an insulator element at the endogenous bithorax locus, as well as in transgenic assays for 4 well-studied insulators^31,32,59,60^, but not more globally at endogenous TAD boundaries. Even for these individual cases, although specific motifs were shown to be essential, the pairing rules that gives rise to their directionality was not elucidated. Our functional assessment of pairing compatibility uncovered heterotypic motif pairs as the predominant signature that explains working boundary pairs. This complexity of motif combinations likely masked their detection previously, and required this prior functional knowledge.

### Specific promoter motifs, but not the presence of a promoter *per se*, are correlated with boundary activity

In the experimentally tested boundaries, promoter presence and their direction are not informative to distinguish working from non-working boundary pairs. Active transcription may therefore have a limited impact on the formation of TAD boundaries, as also suggested by the subtle effects observed after the pharmacological inhibition of transcriptional elongation on TAD structure^30,61^. On the other hand, some promoter motifs are associated with working boundary pairs, suggesting that specific promoter types could contribute to boundary function, probably via the binding of dedicated proteins. M1BP and Trl, for instance, have been reported to bind at promoters at TAD boundaries^30,62^; the M1BP motif is enriched in the working boundary set, albeit at lower frequency than the 54 predictive conserved motif pairs. The motif pairs also include multiple AT-rich homeobox motifs, which are enriched in transcriptionally active regions.

### TAD folding likely requires heterotypic protein-protein interactions

While many *Drosophila* insulator proteins can form homodimers^63^, homodimerization between insulators bound at the two loop anchors is not sufficient to explain *Drosophila* TAD boundaries, at least for the known proteins. Many TAD boundaries are occupied by different combinations of insulator proteins, for example CTCF on one side and BEAF-32 on the other (e.g. Fig. 1G). For example, the boundaries flanking the *ftz* locus (called SF1 and SF2) are bound by CTCF on one side and Pita on the other^64^. Although many of these proteins can homodimerize, including CTCF, Su(Hw) and Zw5, they can also form heterodimers with other insulator proteins. Such heterodimers can form through direct protein-protein interactions, for example BEAF-32 with Z4 and Chromator proteins^65^ or indirectly through co-regulators. For example, almost all known C2H2 insulator proteins can bind to CP190^66^, which could therefore bridge across proteins bound to heterotypic sites. Moreover, many of the newly identified directional motif pairs identified here within working boundary pairs, are bound by proteins known to engage in heterotypic protein-protein interactions, like Bbx with Su(Hw)^67^ and Myb with Pita^68,69^, while others have domains compatible with protein interactions, like the ZAD, BTB/POZ and BESS domains. Similarly, some of these proteins can also bind to PcG domains, such as Myb^55^, Aef-1^70^, CG7878/Mettl14^71^, whose bound complex PRC1 can functionally interact with cohesin in *Drosophila* embryos^72^. Taken together, our results suggest that when these boundary regions come into physical proximity with each-other (by a mechanism yet to be elucidated), the proteins bound to them can interact by direct or indirect protein-protein interactions, likely stabilizing the loop structure in a manner similar to CTCF::CTCF and CTCF::cohesin interactions in mammals^14,73,74^.

## ACKNOWLEDGEMENTS

We thank all members of the Furlong lab for discussions and very useful comments on the manuscript. This work was technically supported by the EMBL Genomics Core (GeneCore) and Advanced Light Microscopy (ALMF) facilities. This work was financially supported in part by an ERC advanced grant DeCRyPT (787611) and DFG-SPP 2202 grant to E.E.F.

## AUTHOR CONTRIBUTIONS

M.V., G.R.C. and E.E.F. designed the study. M.V. and G.R.C. generated all boundary insertion lines, collected embryos, and performed the DNA FISH. R.R.V performed the Tiled-Capture Hi-C, and C.G. made the contact matrices. M.V. performed all other data analysis and motif enrichments, with help from M.F. for SNP calling and A.C. for cross-species sequence alignments. C.S. performed RNAi knockdowns and DNA FISH, analyzed by M.V. M.V., G.R.C.. and E.E.F. wrote the manuscript with input from all authors.

## ONLINE METHODS

### Defining TAD boundaries and their insulator occupancy

To initiate this study, we first defined endogenous TAD boundaries at 6-8h (roughly mid-embryogenesis). Using our previously published deeply sequenced Hi-C data from 6-8h (stage 11/12) embryos^76^, we called TAD boundaries at different resolutions: 1, 2, 5, and 10 kb (Fig. M1). TAD calls by hicFindTADs define consecutive boundaries as a pair, and do not consider multi-way interactions or sub-TADs. To overcome this, boundary calls at different resolutions were manually inspected alongside Hi-C maps using the HiGlass tool^77^ in a Docker container, and the boundary pairs forming a TAD were manually annotated, allowing for TAD nesting and multi-way interactions between boundaries (Fig. S9). All boundaries were resized to 2 kb, leading to a final high-quality set of 1567 boundaries (Table S11). We also performed ChIP-seq on four major insulator proteins (BEAF-32, CTCF, Su(Hw) and CP190) on embryos at the same stage (6-8h), to classify boundaries based on combinatorial insulator binding (BioProject accession number PRJEB38618). The occupancy of these four insulators at the 2kb boundaries is shown in Figure 1B. We used these data to select endogenous boundaries that represent a range of occupancy, promoter features and insulation.

### Generation of TAD boundary insertion lines

The 16 selected TAD boundaries were inserted into TAD A or TAD B by attP-attB recombination-mediated cassette exchange using the MiMIC system^47^. Selected boundaries were PCR-amplified from purified *Drosophila* genomic DNA and inserted into the donor plasmid pBS-KS-attB1-2-DsRed-loxP via InFusion cloning in the PstI site, upstream of the *dsRed* reporter gene flanked by loxP sites. Primers for InFusion cloning were designed using the InFusion cloning tool in SnapGene v4.1.9. The list of oligonucleotides used for TAD boundary cloning can be found in Supplementary Table S12, and the boundary coordinates in Supplementary Table S13.

The donor plasmid was injected into BDGP lines (BDSC #33194 (TAD A), BDSC #60957 (TAD B)) with a MiMIC landing site, the phiC31 integrase and Flp recombinase. All male adults derived from the injected embryos were crossed to *yw* virgins. In the F1, DsRed-positive male transformants were crossed to virgin females with a GFP balancer chromosome. In the F2, positive siblings were crossed, and the F3 screened for GFP-negative homozygous mutants. Homozygous viable stocks were maintained in homozygosity, and homozygous lethal lines were maintained over the GFP balancer.

F1 crosses were genotyped to detect the orientation of the boundary insertion as described in ^47^. The correct landing site and the boundary sequence were verified by Sanger sequencing in the final homozygous fly line, using primers in the genomic region flanking the MiMIC insertion site to amplify the full cassette. Primers used for orientation assessment and genotyping are reported in Supplementary Table S14.

### DNA Fluorescence *In Situ* Hybridization (DNA FISH)

#### Embryo fixation

Adult male and female flies were placed in an embryo collection cage with an apple juice agar plate and yeast paste, at 25°C. After three consecutive 1 h pre-lays, embryos were collected on a new plate for 2 hours and aged at 25°C up to the desired time point (6-8 h). Embryos were then washed with dH2O into a container with a mesh, dechorionated in 8% NaOCl (50% household bleach in H2O) for 2 min, rinsed in dH2O and PBTr 0.1% (PBS 1X + Triton 0.1%) and dried on blotting paper. Dechorionated embryos were fixed in a 1:1 mixture of fixative solution (4% methanol-free paraformaldehyde diluted in PBS 1X) and heptane on a shaker at 470 rpm for 20 min. Then, the lower fixative phase was removed. Devitellinization was performed by adding methanol to the remaining heptane phase and vigorously shaking for 1 min, until most embryos settled at the bottom of the tube. Embryos were finally washed three times in pure methanol and stored in methanol at -20°C.

#### DNA FISH

DNA fluorescent *in situ* hybridization (DNA FISH) probes were designed on TAD boundaries and anchors of high-frequency interaction, as visualized in the Hi-C browser Juicebox^78^. For TAD A, primers for amplification of the selected 8 kb regions from genomic DNA were designed using Primer3 in the Primer Blast tool from NCBI (https://www.ncbi.nlm.nih.gov/tools/primer-blast/), and are reported in Supplementary Table S15. After PCR and gel band extraction, the A-tailed purified DNA was ligated into the linearized pGEMT-Easy vector (#A1360, Promega), according to manufacturer’s instructions. The insertion was confirmed by gel electrophoresis and sequence-verified with T7 and Sp6 standard primers. For TAD B, 80 kb BACs from the BACPAC Resource Center (Supplementary Table S16) were directly used as probes after fluorescent labelling. Probes were fluorescently labelled using the Nick Translation Kit (#7J0001, Abbott Bioscience) and fluorescently labelled dUTP nucleotides (Aminoallyl-dUTP-ATTO-550 #NU-803-550-S, Jena Bioscience; Alexa Fluor 647-aha-dUTP #A32763 Thermo Scientific), according to manufacturer’s instructions. Probes were precipitated separately and only mixed shortly before hybridization. For each embryo sample, 5 µL of fluorescently labelled probe were mixed with 2 µL salmon sperm DNA (#D7656, Sigma Aldrich), 0.1 volume 3 M NaOAc (pH 5.2) and 3 volumes of ethanol, vortexed and precipitated by centrifugation at 17000 g and 4°C for 30 min. After washing in 70% ethanol, the pellet was dried and formamide was added according to the following formula: µL formamide = N × 6.25 µL ÷ n (N = number of embryo samples, n = number of probes per sample). The resuspended probes were left on a shaker at 37°C until prior to hybridization. At that point, probes were denatured at 75°C for 10 min, briefly cooled down on ice, and mixed with 2x hybridization buffer (80% Dextran 50, 4x SSC pH 7.5, 0.055 µg/µL BSA, and 2 µg/µL RNAse A).

Embryos fixed in 4% paraformaldehyde and stored in 100% methanol at -20°C were rehydrated by sequential washes in 75%, 50% and 25% methanol in 2x SSCT (2x SSC pH 7.0 + 0.1% Tween-20) for 5 min, followed by three 5-min washes in 2x SSCT. All washes were performed at RT on a nutator, unless otherwise specified. Embryos were then washed for 10 min in 20% FA/2x SSCT pH 7.4 (20% formamide, 2x SSC pH 7.0, 0.1% Tween-20) and for 10 min in 50% FA/2x SSCT pH 7.4 (50% formamide, 2x SSC pH 7.0, 0.1% Tween-20). Embryos were washed twice for 1 h each in 50% FA/2x SSCT pH 7.4 at 37°C on a nutator; at the end of the 2 h, the buffer was removed and embryos were denatured at 80°C for 15 min in a water bath, placed on ice and mixed with the probes denatured at 75°C for 10 min in the hybridization mix. Samples were overlaid with a drop of mineral oil to prevent evaporation and incubated overnight at 37°C for hybridization. The next day, after removing the mineral oil, samples were washed twice for 30 min in 50% FA/2x SSCT pH 7.4 at 37°C in a thermomixer, and once in 20% FA/2x SSCT pH 7.4 at RT on a nutator for 30 min, followed by three 10-min washes in 2x SSCT. Finally, the supernatant was removed and embryos were mounted in ProLong Gold Antifade Mountant with DAPI (#P36935, Thermo Fisher) on a glass slide, covered with a coverslip and sealed with nail polish.

#### Combined DNA/RNA FISH

The combined DNA/RNA FISH was performed in all TAD B insertion lines. Since some lines were homozygous lethal and had to be maintained over a GFP balancer, RNA FISH was incorporated into the DNA FISH protocol to detect the GFP signal and select GFP-embryos homozygous for the insertion for imaging. For consistency, DNA/RNA FISH was performed on all TAD B lines, including the homozygous viable lines.

The first part of the combined DNA/RNA FISH consists of the DNA FISH protocol described above, except that the hybridization mix contained a RNase inhibitor VRC (2 µM) (#R3380, Sigma Aldrich) instead of the RNAse. After the last SSCT wash, RNA detection was started using the HCR v3.0 kit (Molecular Instruments), following manufacturer’s instructions. After three 5-min washes in PBTw 0.1% (PBS 1X + Tween 0.1%), samples were postfixed in 4% formaldehyde in PBTw 0.1% for 20 min on a nutator, and washed five times for 5 min each in PBTw 0.1%. Samples were then pre-hybridized in 30% probe hybridization buffer for 30 min at 37°C, and incubated overnight at the same temperature with the probes diluted in 30% probe hybridization buffer. The next day, excess probes were removed by four 15-min washes in 30% probe wash buffer at 37°C, and samples were washed three times for 5 min each in 5x SSCT (5x SSC + 0.1% Tween-20). The probe signal was amplified using fluorescently labelled hairpins: following a pre-amplification step in amplification buffer for 30 min on a nutator at RT, samples were incubated with activated fluorescently-labelled hairpins overnight. The next day, excess hairpins were removed by five washes in 5x SSCT at RT on a nutator, to a total of 1 h washing time. Finally, the supernatant was removed and embryos were mounted in ProLong Gold Antifade Mountant with DAPI (#P36935, Thermo Fisher) on a glass slide.

#### Image acquisition

DNA FISH and DNA/RNA FISH samples were imaged on a Leica SP8 confocal microscope equipped with the HC PL Apo CS2 64x/1.4NA Oil and the HC PL Apo CS2 100x/1.4NA Oil objectives, 405 nm laser, white light laser (470-670 nm) and HyD detectors. Over 200 nuclei from 3-4 dorsally oriented stage 11 embryos per line were imaged acquiring a Z-stack with 200 nm step size spanning the entire outermost nuclei layer in the centre of the dorsal epidermis. For TAD B, only GFP-negative homozygous embryos were imaged.

#### Quantification of 3D distance

Images were deconvolved using the Huygens Professional software (SVI) with default parameters. 3D distances between DNA FISH probes were measured using the custom Fiji^79^ plugin *Analyze FISH spots*. The plugin first performs spot detection, and then computes pairwise 3D distances between each spot and their nearest neighbour in the other channels. The coordinates of each spot were determined starting from a manual spot selection using the multi-point tool. Inclusion criteria are that the spot must be present in ≥ 3 consecutive stacks, have the highest intensity in the middle stack and be in an interphase nucleus. At the end of the manual selection of spots on the whole image, the plugin reports the x, y and z coordinates with the maximum intensity in each of the DNA FISH probe channels for each spot, and computes the pairwise distances. Specifically, the plugin examined the Gaussian distribution of the fluorescence intensity of the spot along the x, y and z axis, finding the centre of the spots as the maximum of the Gaussians, and uses the difference of Gaussians (DoG) to calculate 3D distance between spots. Multiple lines were quantified independently by two different experimenters, with highly similar measurements. The distribution of distances between each given condition and the WT was compared by a Kolmogorov-Smirnov test. For data visualization, violin/boxplots were trimmed at 1 µm, but all data points were used for the analysis.

### Multiplexed Tiled Capture Hi-C

#### Embryo collection and fixation for capture Hi-C

Staged embryo collections (as described above) were aged at 25°C up to the desired time point (4-20 h), and fixed with 10 ml cross-linking solution (50 mM Hepes, 1 mM EDTA, 0.5 mM EGTA, 100 mM NaCl, 1.8% formaldehyde, pH 8) in 30 ml heptane, by shaking at 470 rpm for 15 min. After a brief spin at 500 g for 1 min, fixative and heptane were removed, and fixation was quenched in a stopping solution of 125 mM glycine in PBTr 0.1% by vigorously agitating for 1 min. Then, embryos were gently pelleted by centrifuging at 500 g for 1 min, washed twice in PBTr 0.1% and dried on a 125 μm Nitex membrane (#03-125/45, Sefar Nitex Switzerland) over blotting paper. Dried embryos were transferred to a clean tube, weighted, snap-frozen and stored at -70°C until further use.

#### Large scale nuclear isolation for Hi-C

Nuclei were isolated from 0.5–1.2 g of 1.8% formaldehyde-fixed embryos and stored at -70°C. Frozen embryos were thawed in 10 mL cold HB buffer (15 mM Tris-HCl pH 7.4, 15 mM NaCl, 60 mM KCl, 340 mM sucrose, 0.2 mM EDTA pH 8.0, 0.2 mM EGTA pH 8.0, 1x Roche cOmplete Protease inhibitors), and dounced 20 times with a loose pestle and 10 times with a tight pestle in a 15 mL tissue homogenizer (Wheaton). The homogenized suspension was filtered through two perpendicular layers of Miracloth (#475855-1R, Calbiochem) and centrifuged for 10 min at 3200 g at 4°C. After removing the supernatant, the nuclei pellet was resuspended in 10 mL HB buffer and centrifuged for 10 min at 3200 g at 4°C. The supernatant was removed, and the nuclei in the pellet resuspended in 3 mL PBTr 0.1% with 1x Roche cOmplete Protease inhibitors and dissociated by passing them 10 times through a 20G needle and 10 times through a 22G needle using a 5 mL syringe. The suspension was then filtered through a 20 μm Nitex membrane (#03-20/14, Sefar Nitex Switzerland).

Nuclei were counted using CountBright Absolute Counting Beads on BD LSRFortessaTM X-20 Flow Cytometer. Nuclei were spun down at 2000g for 10 min at 4°C, the supernatant was removed, samples were snap-frozen and stored at -70°C).

#### Multiplexed Tiled Capture Hi-C

Tiled Capture Hi-C (cHi-C) libraries were generated using a Bridge-Adaptor *in situ* Hi-C protocol^11^, which leverages biotinylated bridge oligo adaptors for multiplexing between samples^80^, combined to the capture of the genomic regions of interest (ROIs). The captured regions include the two acceptor TADs, extended by ≥ 150 kb on both sides as to cover between 500-950 kb for each TAD, and three control regions (Supplementary Table S17). The set of 13,784 75-bp capture probes, each designed on either end of a DpnII fragment, covered 90% of each region, which were employed for a total panel size of 3.7 Mb. Digestion fragments shorter than the probe length were excluded.

A total of 25 million nuclei stored at -70°C were thawed in 1 mL ice-cold lysis buffer (10 mM Tris-HCl pH 8.0, 10 mM NaCl, 0.2% Igepal CA-630, 1x Roche cOmplete Protease Inhibitors), incubated on ice for 30 min and centrifuged at 1000 g 4°C for 5 min. After removing the supernatant, nuclei were resuspended in 200 µL 0.5% SDS and permeabilized at 65°C for 10 min. SDS was quenched by adding 100 µL of 10% Triton and incubating at 37°C for 15 min. Chromatin was digested overnight at 37°C by adding 8 µL of the restriction enzyme DpnII (#R0543, NEB) in 50 µL of 10x DpnII buffer and 130 µL nuclease-free water. Then, another 5 µL DpnII were added in the morning. After 1 h of incubation, the sample was centrifuged for 5 min at 1000 g at RT and the pellet resuspended in 30 µL water. This step was followed by ligation of biotinylated bridge oligos to the fragmented DNA by adding 1 µL T4 DNA ligase HC (#EL0013, Thermo Fisher Scientific) and 5 µL 90 μM annealed biotinylated oligos (MKVII_F_22: 5’-GATCGGCGCTCAGAA/iBiodT/T-3’ and MKVII_R_22: 5’-CTGAGCGCC-3’) to the resuspended pellet, followed by the addition of 4 µL 10x BSA, 5 µL ligation buffer and 5 µL PEG 4000. The reaction was incubated at 16°C overnight and stopped by adding 2.5 µL 0.5 M EDTA. Nuclei were centrifuged and washed four times in a solution with 1x BSA and decreasing SDS concentration from 0.2% to 0%. The nuclei were then resuspended in 250 µL water containing 1x BSA and 0.2% Triton X-100.

At this point, the biotinylated bridge adaptor oligos were ligated to the DNA fragments but not to each other, due to the lack of a phosphate group at their ends. To enable adaptor::adaptor ligation, nuclei were treated with PNK (#M0201L, NEB) in 1x T4 ligase buffer for 1 h at 37°C, followed by ligation at RT for 4 h. The nuclei were centrifuged at 1000 g, 4°C for 5 min, and resuspended in 500 µL of extraction buffer (50 mM Tris-Cl pH 8.0, 50 mM NaCl, 1 mM EDTA pH 8.0, 1% SDS). Proteins were digested by incubation with 20 µL of 20 mg/mL Proteinase K (Invitrogen) at 55°C, 1000 rpm shaking for 30 min, followed by the addition of 130 µL of 5 M NaCl and decrosslinking on a shaker at 68°C and at 1000 rpm overnight.

The next day, 63 µL of 3 M NaOAc pH 5.2, 2 µL of 15 mg/mL GlycoBlue (Life Technologies) and 1 mL of absolute ethanol were added, and the samples incubated at -70°C for 15 min. The DNA was precipitated by centrifugation for 1 h at 20000 g at 4°C, and the pellet washed, dried and resuspended in 50 µL of 10 mM Tris-HCl pH 8.0. RNA was digested with 0.2 μg/µL RNase A (#10109142001, Roche) at 37°C for 15 min. The DNA was sheared ∼200-400 bp with Diagenode Bioruptor Pico (4°C, 8 cycles, 30”/90” ON/OFF). The DNA was size-selected with AMPure XP beads 1–0.6x according to manufacturer’s instructions, eluted in 100 µL of 10 mM Tris-HCl pH 8.0, and quantified with Qubit. Biotinylated DNA fragments were captured with 30 μL of Dynabeads Streptavidin M-280 beads (#11205D, Thermo Fisher Scientific), followed by washing in bead wash buffer (5mM Tris-HCl pH 7.4, 0.5 mM EDTA pH 8.0, 1M NaCl, 0.1% Tween-20) and resuspension in 40 μL of 10 mM Tris-HCl pH 8.0.

The sequencing library was prepared using the Swift Biosciences Accel-NGS 2S Plus DNA Library Kit (Swift biosciences, 21024), with slight modifications to allow library preparation from DNA bound to Streptavidin beads (from the library preparation published by Arima https://arimagenomics.com/wp-content/files/User-Guide-Library-Preparation-using-Swift-Biosciences-Accel-NGS-2S-Plus-DNA-Library-Kit.pdf). The library was amplified with eight PCR cycles, quantified by Qubit and checked on a Bioanalyzer.

Samples with compatible indexes were pooled and processed according to the Twist Target Enrichment protocol, with minor modifications as follows: The indexed libraries were dried using a vacuum concentrator and resuspended in a blocker solution containing 5 µL SeqCapEZ from NibleGen (#06684335001, Roche) and 7 µL of universal blockers. Then, 1 µL of 10 µM block bridge oligo (block_bridge_22: 5’-GATCGGCGCTCAGAATT[SpcC3]-3’) was added as an additional blocker to block the bridge adapter sequence. Meanwhile, a probe solution was prepared by adding 4 µL of capture Hi-C probes to the Hybridization Mix, heated at 65°C for 10 min and slowly cooled down for 5 min at RT. For hybridization, the probe solution was heated to 95°C for 2 min and cooled on ice for 5 min, the library pool was heated to 95°C for 5 min, and both were equilibrated at RT for 5 min before mixing. Next, 30 µL Hybridization Enhancer was added and the reaction incubated at 70°C for 16 h. The hybridized library was then bound to streptavidin beads, PCR-amplified for 8–9 cycles and purified using DNA Purification beads. After quantification with Qubit and quality control on a Bioanalyzer (displaying a single peak between 350 and 500 bp), the libraries were sent for sequencing on an Illumina NextSeq 500 platform 150 PE at the EMBL Genomics Core Facility, using standard Illumina primers.

The biotinylated sample-specific bridge-adaptor barcode allowed the nuclei to be pooled from multiple samples for *in situ* Hi-C (involving *in situ* chromatin ligation, sonication and Hi-C library preparation), effectively using the same amount of chromatin as standard Hi-C. Sample pooling, enabled by early sample barcoding, minimizes loss of material from individual samples and reduces inter-sample variability, thereby enhancing comparability among Hi-C libraries, reducing technical bias and cost. Here, we captured and sequenced pools of 6-to-8 samples from 4-20 h embryos for each of the 38 lines.

### Whole genome sequencing (WGS) of TAD B parental line

A number of insetions in TAD B were lethal (14/26), as it contains three essential developemtnal genes. We therefore maintained the insertions over a GFP-marked balancer chromosome. Embryos of mixed genotypes were collected and used for the cHi-C, which includes embryos with homozygous insertion (25%), heterozygous insertion::balancer chromosomes (50%), and homozygous balancer embryos (25%). To distinguish between these genotypes to generate cHi-C matrices for the homozygous boundary insertion lines, we separted all Hi-C reads based on informative SNPs into balancer containing (which were disgarded) and non-balancer reads (used to make the contact matrices). To enable this, we first sequenced the partental TADB insertion line (the MiMIC line BDSC#60957)^81^ before the insertions. 50 adult flies grown at 25°C were snap-frozen in liquid nitrogen. Flies were ground on dry ice in an Eppendorf tube with a pestle. Ground flies were resuspended in 250 µl of Buffer A (100 mM Tris-HCl pH7.5, 100 mM EDTA, 100 mM NaCl, 1% SDS) with 5 µl of proteinase K 20 µg/µl. The sample was incubated at 65°C for 3 hours and at 95°C for 10 min, then spun for 2 min at room temperature. The supernatant was transferred to a new tube and incubated at 37°C for 45 min with 2 µl of RNaseA 100 mg/ml. The DNA was extracted by two rounds of standard phenol-chloroform extraction, precipitated with ethanol, and resuspended in 150 µl of Tris-HCl 10 mM. 500 ng DNA were sonicated in 150 μl Tris-HCl 10 mM (5 pulses, 30’’ on 30’’ off) to obtain 200-400 bp fragments. DNA was size-selected with AMPure beads 1.4x. Library preparation was done with the NEBNext Ultra II DNA Library Prep kit according to manufacturer’s instructions.

### RNAi depletion of candidate TFs

Maternal and early zygotic depletion of candidate TFs was performed by RNAi-mediated knockdown, utilizing stocks that carry short hairpin RNAs (shRNAs) under the control of UAS elements as described previously^82^. Virgin females of the respective shRNA line [obtained from the Bloomington *Drosophila* Stock Center (BDSC): Bbx (BDSC, #57553); Sox21b (BDSC, #60120); Myb (BDSC, #35053); Pita (BDSC, #57732); CG4404 (BDSC, #62415); CG12155 (BDSC, #26232)] were crossed to males from the Maternal-tubulin-GAL4 line (BDSC, #7063), which expresses GAL4 during late stages of oogenesis^82^ and incubated at 25°. To obtain embryos depleted of maternal transcript of each factor, F1 virgin females carrying one copy of the UAS-shRNAs and one copy of the Maternal-tubulin-GAL4 transgene were backcrossed to males carrying the UAS-shRNAs construct at 25°. From these crosses, F2 embryos at 2-4 h after egg laying were collected at 25° for DNA FISH experiments. Embryo fixation and DNA FISH were performed as described above.

## Computational Methods

### Capture Hi-C data processing

#### Read separation

As homozygous lethal insertions into TAD B were maintained over a GFP balancer, embryo collections are a mixture of 25% homozygous ‘boundary insertion’ embryos, 25% homozygous balancer embryos and 50% heterozygous embryos with the insertion over the balancer. To examine the effect of the insertion without confounders from the balancer signal, reads were separated by genotype based on the GATK Best Practice Workflow in the MiMIC and in the balancer chromosome (inferred from a virginizer|Balancer F1 cross and Virginizer line, as in ^48^), using a custom script^83^.

The parental genomic DNA from three parental lines (MiMIC, virginizer and virginizer|Balancer cross) were trimmed using trim_galore (v. 0.5.0)^84^ and options “-q 30 - phred33 --fastqc --illumina --length 25 --paired” and mapped to the joint reference genome of *D. melanogaster* (dm6) and *Wolbachia* (AE017196) with bwa mem (v. 0.7.17)^85^ and option “-T 20”. Duplicates were annotated with picard (v. 2.16.0)^86^ MarkDuplicates and reads were filtered with samtools (v. 1.9) and options “-F 1548 -f3”. The resulting bam files were then annotated with picard AddOrReplaceReadGroups to create one group per parental line. Before proceeding with variant call, we performed a base recalibration with GATK (v. 4.1.0.0)^87^ BaseRecalibrator and ApplyBQSR using the DGRP vcf as reference^88^ for common variants in *Drosophila* populations.

Variants were called with GATK HaplotypeCaller^87^ using a joint genotyping strategy and options “-G StandardAnnotation --min-base-quality-score=20”. Variants on balancer chromosomes were inferred from the virginizer|Balancer F1 cross and Virginizer line as follows: (1) biallelic heterozygous variants in the virginizer|Balancer F1 cross genotyped as homozygous in the Virginizer line were genotyped as homozygous for the oppositive allele in the Balancer line; (2) biallelic variant homozygous for the same allele in both virginizer|Balancer F1 cross and Virginizer line were annotated for the same genotype in the Balancer chromosome; (3) all other variants were annotated as unknown genotype for the Balancer chromosome. Next, variants were filtered applying a set of hard cut-offs using bcftools (v. 1.9)^89^: “MQ > 58 & MQRankSum > -2.5 & MQRankSum < 2.5 & QD > 15 & SOR < 1.5 & FS < 10 & ReadPosRankSum > -6 & ReadPosRankSum < 6”.

Capture-Hi-C reads from F1 crosses were then mapped on the *D. melanogaster* dm6 assembly with BWA-MEM^85^ with default parameters. The resulting bam files were then split in three .bam files corresponding to MiMIC, balancer and unknown genotypes with a custom script. Reads were piled up on variants obtained from the previous step. Reads unambiguously harbouring alleles from MiMIC or balancer genotype were assigned to the corresponding genotype, while reads that did not overlap known variants or harboured ambiguous alleles were assigned to the unknown .bam file. The original fastq files were then filtered for reads belonging to the split bam files with seqtk (v. 1.3) to produce a MiMiC and a Balancer fastq file for each sample.

Since trans interactions were not used for subsequent analysis, all data from the allele with the insertion (i.e. from the homozygous insertion and heterozygous embryos) was retrieved. Reads from the insertion allele were processed as described in the next section.

#### Capture Hi-C data processing

The Illumina sequencing files were demultiplexed, converted into FASTQ format and quality-checked using FastQC^90^ in Galaxy^91^ Reads were mapped on the *D. melanogaster* dm6 genome assembly using BWA-MEM^85^ with default parameters (Galaxy Tool version 0.7.17.1). Hi-C matrices were generated using the HiCExplorer^23^. BAM files were sorted by name using the Galaxy tool “Sort reads by name using Samtools sort” (v. 2.0.2)^89^ and processed using hicBuildMatrix (Galaxy tool version 3.6+galaxy0) with the list of DpnII restriction sites as produced by hicFindRestSite (Galaxy version 3.6+galaxy0, with --searchPattern GATC) ran on the dm6 FASTA genome and options –restrictionSequence GATC --danglingSequence GATC --binSize 1000 – skipDuplicationCheck to yield a matrix with 1 kb resolution; 2 and 5 kb resolution matrices were also computed. Quality control was performed on the initial complete raw data using the quality reports produced by hicBuildMatrix and the MultiQC tool^92^, to obtain the number of duplicated read pairs, dangling end, self-ligations, the distribution by read orientation and the proportion of short-range, long-range and trans-chromosomal interactions.

The subsequent operations were performed using custom scripts and HiCExplorer v. 3.7.2. Trans reads were removed by hicAdjustMatrix, keeping only chromosomes 2L, 2R, 3L, 3R, 4 and X (options -a keep --interIntraHandling inter --chromosomes chr2L chr2R chr3L chr3R chr4 chrX). Technical replicates were merged by hicSumMatrices and converted from hic to cool format using hicConvertFormat. To avoid bias from the interactions with only one end in the captured region during the normalization step, the matrix was subset using hicAdjustMatrix to include only the contacts with both ends within the captured region. Each line with an insertion was compared to the wildtype (WT) starting line (without the insertion) of the same TAD separately: the total reads in each condition were computed, and the matrix of the sample with the highest coverage was downsampled to the reads of the sample with the lowest read number using cooltools random-sample (v. 0.4.0). Both the WT and the insertion line capture Hi-C matrices were then Knight-Ruiz(KR)-normalized using hicCorrectMatrix with the – correctionMethod KR parameter^93^.

Acceptor TADs were visualized using hicPlotTADs (in HiCExplorer v. 3.7.2)^23^ on subsampled and KR-normalized capture Hi-C matrices at 5 kb resolution, including only the captured ROI.

### Identification of boundary pairs by virtual 4C from Capture Hi-C data

The boundary pairs formed by TAD boundary insertions were visualized in hicPlotTADs using the capture Hi-C matrices to generate virtual 4C profiles. The effect of the insertion was evaluated based on the capture Hi-C matrix, and on the subtraction of the contact frequency in the insertion versus the WT using hicCompareMatrices (in HiCExplorer v. 3.7.2)^23^. The result was confirmed by computing the insulation score at 5 kb resolution at the insertion site with hicFindTADs and FDR-adjusted p value < 0.01.

To understand which acceptor TAD boundaries interact with the inserted boundary, virtual 4C plots were generated from capture Hi-C data with hicPlotViewpoint (in HiCExplorer v. 3.7.2)^23^, using the MiMIC insertion site or the location of DNA FISH probes (left and right anchor in TAD A, left and right boundary in TAD B) as viewpoint. Interactions were computed in the pairwise normalized WT and insertion lines, and the contacts made by the insertion were detected as a gain of interactions between the acceptor TAD boundaries and the insertion site (Table S3).

### Functional conservation of inserted boundaries

To assess if the function of the 16 inserted boundaries is conserved across the *D. melanogaster* clade we evaluated TAD syntenic blocks^50^ in 11 species (*D. ananassae*, *D. elegans*, *D. triauraria*, *D. biarmipes*, *D. takahashii*, *D. eugracilis*, *D. erecta*, *D. yakuba*, *D. simulans*, *D. ficusphila*, and *D. melanogaster*) with at least 25 Mya divergence (Supplementary Table 1 from ^50^). Conserved boundaries were defined as boundaries between the same two orthologous TADs in multiple species, or within 1 Hi-C bin (5 kb) from an orthologous TAD.

### Evolutionary conservation of motifs in TAD boundaries

#### Alignment of inserted and acceptor TAD boundaries across the D. melanogaster clade

The evolutionary conservation of inserted and acceptor TAD boundaries was evaluated by aligning their sequence in *D. melanogaster* to the genome of 18 other *Drosophila* species in the same clade. The conservation score of each base was defined as

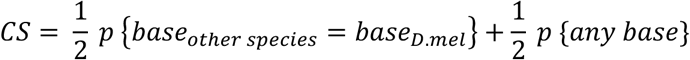

Where the first component is the frequency at which the other species have the same base as in *D. melanogaster*, and the second component is the frequency of any other base (i.e. not having a gap in the alignment). From the alignment, the sequences were mapped back to the *D. melanogaster* genome. To find conserved regulatory regions, we considered the non-coding sequences in Dnase hypersensitive sites (DHSs)^52^ extended by +/-100 bp. The conservation score for each position of the alignment was defined as the mean of the probability of having the same base as *D. melanogaster* and that of having any base in that position. For each boundary the bases with a conservation score in the top 75% of the conservation score were defined as conserved. Consecutive conserved bases were considered a conserved sequence; up to three consecutive nucleotides with lower conservation were allowed within a conserved sequence. Conserved sequences with length < 6 bp were discarded.

#### Motif search and annotation in aligned boundaries

The conserved sequences within the non-coding DHSs were scanned for known motifs using FIMO (MEME suite v. 5.5.5)^53^, with a background model based on nucleotide frequency in the DHSs from Reddington *et al.* 2020 and q value < 0.8. A conserved sequence was assigned to a known motif if it was found in the DHSs and overlapped with the sequence. For the motif search, three different motif sources were used: CISBP 2.00 *D. melanogaster*^94^ for TFs in general, a collection of insulator protein motifs obtained by MEME-ChIP on ChIP-seq data from our lab (for BEAF-32, CTCF, Su(Hw), Trl, M1BP)^29,46^ and other labs (Ibf1, Ibf2, Pita, Zw5, ZIPIC)^95,96^, and core promoter motifs^97,98^. Conserved sequences were classified based on the contained motifs (PWMs) into insulators, other TF and promoter motifs, and sequences with multiple motifs were assigned to mixed categories (e.g. insulator + promoter). Sequences not assigned to any known motif were defined as new motifs. The known and new conserved motifs were used for downstream analysis.

The assignment of *de novo* discovered motifs to proteins is always challenging and depends on available motif databases, and suffers from the limitations in how those resources were constructed (using SELEX, protein-binding arrays, bacterial one hybrid, ChIP-seq). We therefore note that although the motif is enriched, the protein bound may not correspond to the assigned factor in cisBP. Moreover, many discovered motifs are very similar it each other, and it is often not clear if they represent the same motif bound by the same factor, or different variants that are bound by different TFs with similar DNA binding domains. Given this, here we preferred to be conservative and report all pairs as discovered rather than trying to match similar motifs. This has the overall effect to inflate the number of motif pairs, such that the number of unique non-overlapping directional motifs will be lower than 54 pairs, and as a consequence the frequency (or potential importance) of each pair at boundaries would increase.

### Finding conserved motifs linked to working boundaries in the experimental and genome-wide datasets

FIMO (MEME suite v. 5.5.5)^53^ was run on the DHSs extended by +/-100 bp within the experimental inserted boundaries, using all DHS from^52^ as background, a q value < 0.8 and the known and new motifs in the conserved sequences as motif dataset. The position-weight matrices (PWMs) of known motifs were retrieved from the original motif database, and those for new motifs were generated from the evolutionary conservation, using the frequency of each base in the species alignment from section 0, and rescaled to 1. FIMO (MEME suite v. 5.5.5)^53^ was run on genome-wide boundaries using the same parameters.

Motifs were tested for enrichment in TAD boundaries genome-wide using AME from MEME suite v. 5.5.5, with parameters --scoring max and --method fisher, using a DHS background from^52^. Only motifs enriched in TAD boundaries with adjusted p value < 0.05 were used for further analysis.

To identify motifs associated with working (interacting) boundaries in the experiments, the percentage of working boundaries with each motif was computed separately for each TAD. Working boundaries were defined as boundaries with an effect in DNA FISH and capture Hi-C; in case of discrepancy (3/37 lines), capture Hi-C was used as reference, as it is more sensitive to subtle changes.

To examine the mechanism of orientation- and context-specific boundary function, we searched for pairs of two motifs, each on one anchor of an experimentally tested boundary pair, in all the possible reciprocal orientations (convergent, divergent, tandem plus or tandem minus). The frequency of interacting boundary pairs among those containing the motif pair in a specific orientation was computed separately for each TAD. For genome-wide boundary pairs, the frequency of boundaries with the motif pair in a specific orientation over all possible orientations was computed to derive the orientation preference of a motif pair. A one-tailed Fisher test was used to assess orientation preference. The results were visualized with the ComplexHeatmap R package v 2.14.0^99,100^. Due to the difference in baseline proportion of working boundary pairs in TAD A (32%) and TAD B (80%), oriented motif pairs always associated to working boundaries in TAD B, and in 80% of the cases in TAD A, were considered as motifs potentially driving successful pairing.

For a preliminary evaluation of the sensitivity and specificity of this set of motif pairs, we computed the percentage of working and non-working experimental pairs with the motif pair set. To evaluate if these motif pairs could explain boundaries in the whole genome as well, their frequency in genome-wide boundary pairs was calculated.

Combinations of more than 2 motifs were evaluated in experimental boundaries by the frequency of co-occurrence of two motif pairs in the same working boundary pair. Motif pairs co-occurring > 70% of the times and not composed of highly similar sequences were considered co-occurring (Table S9), and pairs with 50–70% co-occurrence were considered to be potentially facilitating each other’s function (Table S9).

### Logistic regression

A logistic regression model with elasticnet regularization was trained to classify functional and non-functional boundary pairs using one-hot encoded oriented motif pairs as input features. Scikit-learn v1.5^101^ was used for model training and prediction. The training set included 79 working and 108 non-working boundary pairs from experimentally inserted boundaries (Supplementary Tables S3, S18). The test set comprised 1852 working genome-wide boundary pairs (Supplementary Table S19) and a matched negative set of paired 2 kb sequences with similar DHS and insulator protein profiles (BEAF-32, CTCF, CP190, Su(Hw)) but no boundary activity (insulation score > -0.1 and not called as TAD boundaries) (Supplementary Table S20). The 54 oriented motif pairs associated to working boundary pairs both in TAD A (probability of working >= 80%) and in TAD B (100% working) were used as features. Features were preprocessed to combine highly correlated features into one and to exclude features with zero variance. Model parameters were fine-tuned using LogisticRegressionCV with 5-fold cross-validation, elasticnet penalty, saga solver, balanced accuracy scoring and l1 ratios between 0 and 1 with 0.1 step size. The best performing model was tested on the genome-wide boundary pairs.

Feature selection was performed Recursive Feature Elimination Cross-Validation (RFECV) in scikit-learn combined with a logistic regression estimator, with elasticnet penalty, l1_ratio = 0.01 and saga solver. Balanced accuracy on the inserted and on the genome-wide pairs was evaluated for each model. The ROC curve was displayed using the scikit-learn function RocCurveDisplay.from_estimator.

Since 53% of the genome-wide boundary pairs do not have motif pairs in our set, the theoretical maximum accuracy of the model is 0.73 (i.e., if all the sequence pairs that have a motif pair are predicted to pair, and all those without a motif pair are predicted not to pair). The accuracy of our model, trained exclusively on inserted boundary pairs, is 0.68 on the genome-wide sequence pairs. The use of a test set completely separated from the training set prevents leakage and provides a conservative estimation.

